# Spen and Nito prevent dedifferentiation of progenitors by translationally repressing E(Spl)mγ

**DOI:** 10.1101/2025.08.19.671041

**Authors:** Xiaosu Li, Wenwen Lu, Marisa Connell, Xiaobing Deng, Sijun Zhu

## Abstract

Rapid termination of Notch signaling by asymmetrically segregated Numb is essential for specification of differentiating progeny during asymmetric stem cell division. Using *Drosophila* type II neuroblasts as a model, here we report that specification of the differentiating progeny also requires the expression of the Notch target E(Spl)mγ in the stem cell to be kept at low levels by the SPEN family proteins, Spen and Nito. We show that loss of Spen or Nito leads to a drastic increase in E(Spl)mγ expression in the stem cell and subsequent dedifferentiation of its progeny. Genetic and biochemical studies demonstrate that Spen and Nito maintain the expression of E(Spl)mγ in the stem cell at low levels by directly binding to two identical novel motifs in the 5’UTR to repress its translation. The low expression in the stem cell prevents excessive inheritance of E(Spl)mγ proteins by the differentiating progeny and ensures its rapid removal from the progeny. Together, our work uncovers a novel post-transcriptional mechanism regulating Notch signaling that is critical for cell fate specification during asymmetric stem cell division.

## Introduction

Asymmetric division of stem cells leads to production of two daughter cells with distinct cell fates: a new stem cell and a differentiating progeny. One critical mechanism underlying the acquisition of distinct cell fates of the daughter cells is the asymmetric activation of Notch signaling (Artavanis-Tsakonas et al. 1999). Notch signaling is activated in the newly generated stem cell to promote self-renewal but must be rapidly turned off in the differentiating progeny to allow proper cell fate specification (Le Borgne and Schweisguth 2003; Bhat 2014). Deficiency in Notch signaling can lead to premature depletion of stem cells and developmental defects, whereas aberrant/ectopic activation of Notch signaling may cause the differentiating daughter cell to adopt the stem cell fate and initiate tumor growth (Capobianco et al. 1997; de la Pompa et al. 1997; Henrique et al. 1997; Artavanis-Tsakonas et al. 1999; Hitoshi et al. 2002; Fre et al. 2005; Stockhausen et al. 2010). It has been well established that asymmetrically segregated Numb inactivates Notch in differentiating daughter cells by promoting degradation of Notch, thus terminating the transcription of Notch target genes (Guo et al. 1996; Spana and Doe 1996; Berdnik et al. 2002; Couturier et al. 2012). However, during the asymmetric division, proteins and mRNAs of Notch targets are segregated symmetrically (Fig.1A)(Dong et al. 2012; Liu et al. 2017). Rapid degradation of Notch target mRNAs and proteins is also critical for proper cell fate specification of the differentiating daughter cell, but the underlying mechanisms ensuring the rapid removal of Notch targets are not well understood.

**Figure 1.**
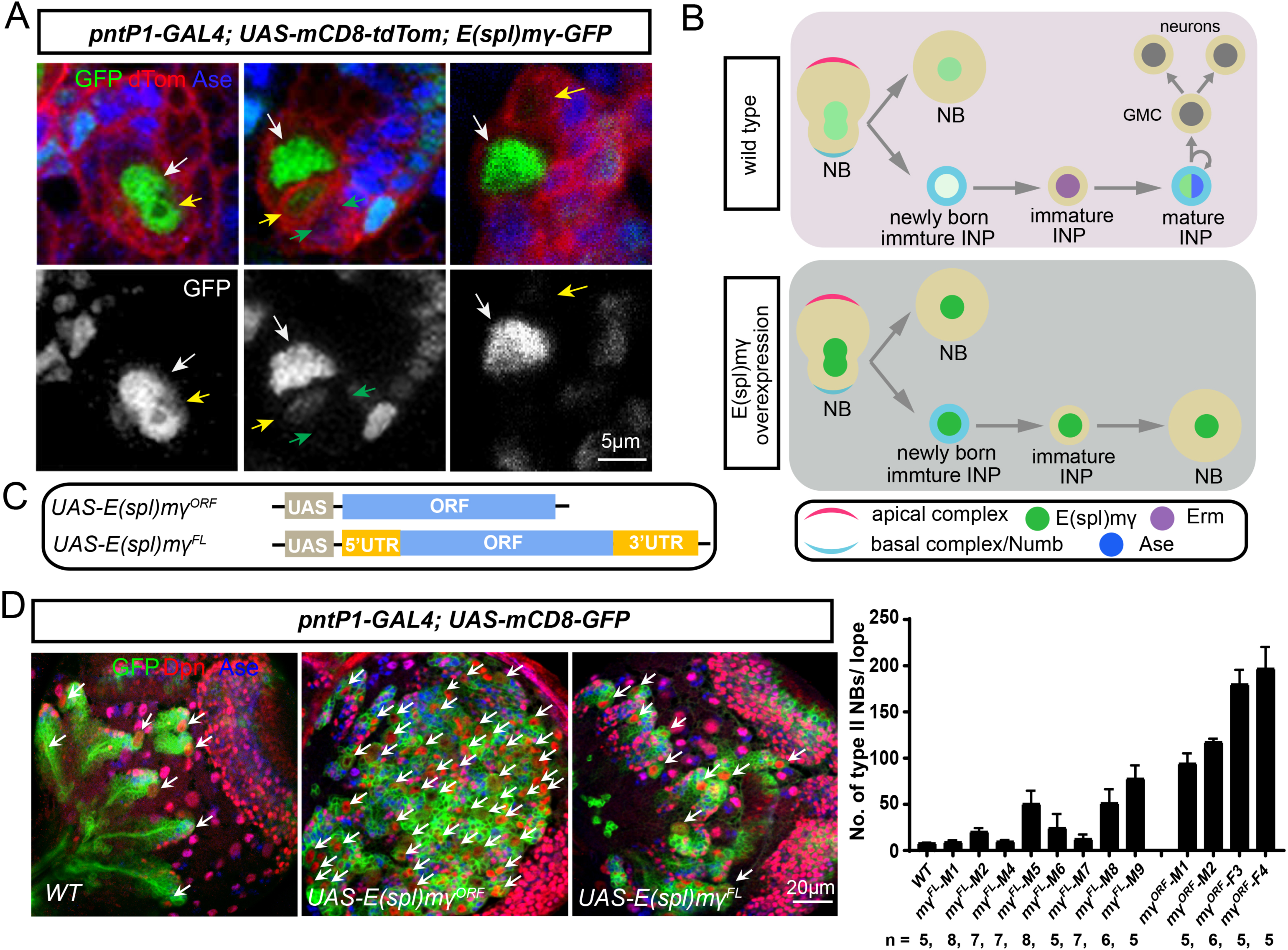
Overexpression of the full-length cDNA of E(Spl)mγ leads to much weaker supernumerary type II NBs phenotypes than the open reading frame alone. (A) E(Spl)mγ-GFP proteins are symmetrically segregated during the division of type II NBs (the left panels) but rapidly degraded in newly generated imINPs (middle and right panels). Type II NB lineages are labeled with mCD8-RFP driven by *pntP1-GAL4* and brains are counterstained with anti-Ase antibody. White arrows point to E(Spl)mγ-GFP in type II NBs; yellow arrows point to E(Spl)mγ-GFP in newly generated imINPs and green arrows to old Ase^-^ imINPs. (B) A diagram of the development of type II NB lineages in wild type and E(Spl)mγ over-expressing brains. In the wild type (upper), E(Spl)mγ expression is quickly lost in the newly generated imINP, which allows PntP1 to activate the expression of Erm in imINPs. Erm in turn inhibits PntP1 expression in imINPs, allowing activation of Ase expression and maturation of INPs. When E(Spl)mγ is overexpressed, E(Spl)mγ inhibits the activation of Erm by PntP1, leading to dedifferentiation imINPs into type II NBs. (C) A diagram of the constructs of *UAS-E(Spl)mγ^ORF^* and *UAS-E(Spl)mγ^FL^*. (D) Compared to a wild type 3^rd^ larval brain lobe that contains only 8 type II NBs (left panel), expression of *UAS-E(Spl)mγ^ORF^*dramatically increases the number of type II NBs (middle panel), but expression of *UAS-E(spl)mγ^FL^* only mildly increases the number of type II NBs (right panel). Type II NB lineages are labeled with mCD8-GFP driven by *pntP1-GAL4* and brains are counterstained with anti-Dpn and anti-Ase antibodies. Arrows point to type II NBs. Quantifications of the number of type II NBs in brains with indicated genotypes are shown in the bar graph to the right.

In *Drosophila* type II neuroblast (NB) lineages, NBs divide asymmetrically to produce a self-renewing NB and an immature intermediate progenitor (imINP), which differentiate to become mature INPs. INPs then undergo several rounds of self-renewing asymmetric divisions to generate ganglion mother cells (GMCs), which further divide to produce neurons or glial cells (Bello et al. 2008; Boone and Doe 2008; Bowman et al. 2008). Like in other stem cells, Notch signaling is asymmetrically activated in newly generated type II NBs and imINPs after the asymmetric division. In the type II NBs, Notch signaling is activated to maintain type II NB identity and self-renewal by activating Hes family proteins, such as E(Spl)mγ (Bowman et al. 2008; Song and Lu 2012; Zhu et al. 2012; Li et al. 2016; Janssens et al. 2017). In newly generated imINPs, asymmetrically segregated Numb promotes endocytosis and degradation of Notch, which is further facilitated by the retromer complex-mediated trafficking (Li et al. 2018). Meanwhile, the Notch target E(Spl)mγ proteins are symmetrically segregated during asymmetric division of type II NBs but rapidly diminish in imINPs (Liu et al. 2017). In type II NBs, E(Spl)mγ prevent the activation of the expression of the forebrain embryonic zinc finger (Fezf) family protein Earmuff (Erm) by the Ets family protein Pointed P1 (PntP1) in type II NBs (Li et al. 2016; Janssens et al. 2017; Li et al. 2017). PntP1 is both necessary and sufficient for the specification of type II NB (Zhu et al. 2011). PntP1 specified type II NBs by inhibiting the expression of the proneural protein Asense (Ase) (Hakes and Brand 2020; Rives-Quinto et al. 2020; Chen et al. 2022). In imINPs, the termination of Notch signaling and rapid removal of E(Spl)mγ allows PntP1 to activate Erm, which in turn promotes INP maturation by interacting with the chromatin remodeling Brahma complex and Histone deacetylase 3 (HDAC3) and providing negative feedback to inhibit the activity and expression of PntP1 in INPs (Weng et al. 2010; Eroglu et al. 2014; Koe et al. 2014; Li et al. 2016; Janssens et al. 2017). When Notch signaling is ectopically activated in imINPs due to either absence of Numb or overexpression of Notch or E(Spl)mγ, imINPs dedifferentiate into type II NBs due to inhibition of Erm expression, leading to supernumerary type II NBs (Fig. 1B) (Wang et al. 2006; Bowman et al. 2008; Li et al. 2017).

Gene expression can be regulated by both transcriptional mechanisms and post-transcriptional mechanisms. Transcriptional regulation of Notch target expression has been well studied and is well conserved among different animal species. Upon binding with the ligand, Notch is cleaved and its intracellular domain translocates to the nucleus, where it forms a complex with Su(H)/RBPJ and co-activator Mastermind (Mam) to activates the transcription of its target genes (Bray 2006; McLaren and Butts 2025). In contrast, post-transcriptional regulation of Notch target expression is less well understood. The untranslated regions (UTRs) at both the 3’ and 5’ end of mRNAs play vital roles in post-transcriptional regulation. UTRs contain cis-acting elements that interact with specific RNA-binding proteins, long non-coding RNAs, and/or microRNAs. UTRs mediate the regulation of stability, translation, and localization of mRNAs, and can exert positive or negative effects on gene expression (Mignone et al. 2002; Hardy and Balcerowicz 2024). E(Spl)mγ transcripts contain a 93-bp 5’UTR and a 201-bp 3’UTR (Artavanis-Tsakonas et al. 1999; Rubin et al. 2000). Previous studies with artificial GFP or lacZ reporter assays showed that the 3’-UTRs of *E(spl)* genes harbor binding sites for multiple classes of miRNAs and mediate the suppression by miRNAs (Lai and Posakony 1997; Lai et al. 1998; Lai et al. 2005). In vertebrates, microRNAi-9 negatively regulates Hes1 expression and contributes to the oscillation of Hes1 expression (Bonev et al. 2012). However, whether the UTRs have any role in regulating the expression of endogenous E(Spl)mγ has rarely been investigated.

Spen (Split End) and Nito (Spenito) belong to the evolutionarily conserved SPEN protein family, which contains three N-terminal RNA-recognition motifs (RRMs) and a C-terminal SPOC (Spen Paralog and Ortholog C-terminus) domain, but Nito is much smaller than Spen (Wiellette et al. 1999; Sanchez-Pulido et al. 2004; Jemc and Rebay 2006). SPEN family proteins play important roles in many developmental processes, such as cell fate specification, axon extension, X-chromosome inactivation, sex-determination, intestinal and germ line stem cell self-renewal (Chen and Rebay 2000; Kuang et al. 2000; Doroquez et al. 2007; McHugh et al. 2015; Querenet et al. 2015; Yan and Perrimon 2015; Andriatsilavo et al. 2018). In humans, Spen (or SHARP), and Nito (or RBM15) are frequently mutated in a variety of cancers, such as leukemia, adenoid cystic carcinoma, bladder cancer, and glioblastoma, indicating that they could function as tumor suppressors (Ma et al. 2001; Frierson and Moskaluk 2013; van der Heijden et al. 2016; Liu et al. 2018; Naderi 2018). Depending on the context, Spen and Nito could act antagonistically or additively/synergistically to regulate various signaling pathways (Chen and Rebay 2000; Kuang et al. 2000; Doroquez et al. 2007; Ma et al. 2007; Chang et al. 2008). In mammals, Spen/SHARP has been shown to act as a transcriptional repressor to inhibit Notch target expression by recruiting HDAC1 repressor complexes (Oswald et al. 2002).

In this study, we investigated post-transcriptional regulation of E(Spl)mγ expression in type II NB lineages. We show here that both the 5’- and 3’-UTRs are required to maintain the expression of E(Spl)mγ at low levels in type II NBs and imINPs. By screening RNA-binding proteins by RNAi, we found that Spen and Nito function partial redundantly to prevent dedifferentiation of imINPs by keeping the expression of E(Spl)mγ in type II NBs at low levels. However, unlike mammalian Spen/SHARP that functions as a transcriptional repressor to inhibit Notch target expression, we demonstrate that *Drosophila* Spen and Nito function as translational repressor to inhibit E(Spl)mγ expression by binding to two identical novel motifs in the 5’UTR. The relative low expression of E(Spl)mγ in type II NBs likely ensures that the amount of E(Spl)mγ proteins segregated into imINPs will not overwhelm the degradation mechanisms, thus allowing rapid removal of E(Spl)mγ from imINPs and maturation of imINPs. Our work demonstrates that in addition to Numb-mediated degradation of Notch, INP fate commitment also requires the expression of E(Spl)mγ in type II NBs to be maintained at low levels by Spen/Nito-mediated translational repression through the 5’UTR. Further, our work reveals a novel mechanism of the regulation of Notch signaling by Spen/Nito.

## RESULTS

### Over-expression of E(spl)mγ with its full-length cDNA leads to much weaker supernumerary type II NB phenotypes than with its open reading frame alone

To investigate if the untranslated regions (UTRs) have any roles in controlling E(spl)mγ expression and INP cell fate specification, we compared overexpression phenotypes of *UAS-E(spl)mγ^ORF^,* which expresses the open reading frame alone of E(spl)mγ, and *UAS-E(spl)mγ^FL^*, which expresses the full-length cDNA of E(spl)mγ including both the 5’-UTR and 3’-UTR (Fig. 1C), in type II NB lineages using *pntP1-GAL4* as a driver. *pntP1-GAL4* is expressed in both type II NBs and imINPs (Zhu et al. 2011). The expression of *UAS-E(spl)mγ^ORF^* using 4 independent lines consistently led to strong supernumerary type II NB phenotypes (with about 100-200 type II NBs/brain lobe on average, compared to 8 per brain lobe in the wild type). In contrast, the expression of *UAS-E(spl)mγ^FL^* led to much weaker phenotypes. Of 8 independent lines we generated, 5 lines gave rise to only 9-25 type II NBs/brain lobe, while the remaining three gave rise to 50-80 type NBs/lobe (Fig. 1D, and Fig. S1). Our previous studies as well as others have shown that the supernumerary type II NB phenotype induced by E(spl)mγ overexpression is due to inhibition of *erm* expression in imINPs and subsequent dedifferentiation of imINPs back to type II NBs (Janssens et al. 2017; Li et al. 2017). The much milder phenotypes resulting from the expression of *UAS-E(spl)mγ^FL^* suggests that inclusion of UTRs leads to lower expression of E(spl)mγ in imINPs. Therefore, the UTRs are likely critical in regulating E(spl)mγ expression and INP cell fate specification.

### The UTRs mediate post-transcriptional repression of E(spl)mγ expression in type II NB lineages

To determine if the UTRs regulate E(spl)mγ expression, we generated nuclear GFP (nGFP) reporter constructs, which utilize the UAS promoter to drive the expression of nGFP coding sequence alone (*UAS-nGFP*), or nGFP coding sequence fused with the 5’UTR (*UAS-5’UTR-nGFP*), the 3’UTR (*UAS-nGFP-3’UTR*), or both (*UAS-5’UTR-nGFP-3’UTR*) (Fig. 2A). These constructs were all inserted at the same attP80 site on the 3^rd^ chromosome to ensure that their expression is not affected by different genomic environments. We expressed these transgenes in type II NB lineages using the *pntP1-GAL4* driver and compared nGFP expression levels in type II NBs and imINPs in age-matched live third instar larval brains. Compared to *UAS-nGFP*, *UAS-5’UTR-nGFP*, *UAS-nGFP-3’UTR*, or *UAS-5’UTR-nGFP-3’UTR* all expressed much lower nGFP levels in both type II NBs and imINPs, with the *UAS-nGFP-3’UTR* giving the lowest nGFP expression (Fig. 2B). These results suggest that both the 5’UTR and 3’UTR mediate post-transcriptional repression of E(spl)mγ expression, but the 3’UTR has a stronger effect. However, the 5’UTR is likely also required to maintain the expression at relatively higher levels given that *UAS-5’UTR-nGFP-3’UTR* gave higher nGFP expression than *UAS-nGFP-3’UTR*.

**Figure 2.**
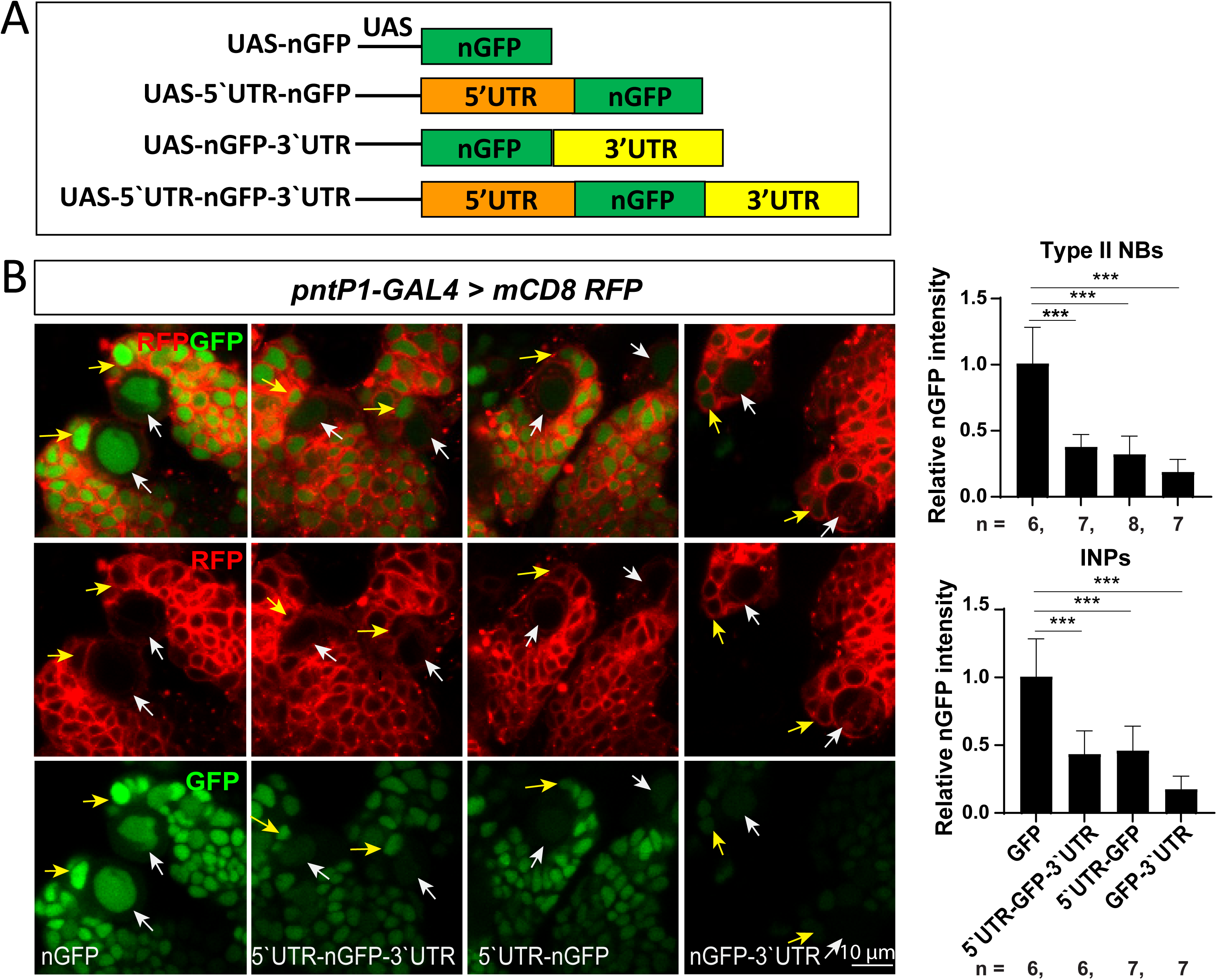
Both the 5’UTR and 3’UTR negatively regulate nGFP reporter expression. (A) Schematic diagrams of the UAS-nGFP reporter constructs without or with the 5’UTR, 3’UTR, or both from E(Spl)mγ. (B) Live images of nGFP expression in type II NB lineages that express *UAS-nGFP*, *UAS-5’UTR-nGFP*, *UAS-nGFP-3’UTR*, or *UAS-5’UTR-nGFP-3’UTR* driven by *pntP1-GAL4*. Type II NB lineages are labeled with mCD8-RFP driven by *pntP1-GAL4*. White arrows point to type II NBs and yellow arrows newly generated imINPs. Bar graphs to the right show quantifications of relative nGFP expression levels in the type II NBs (upper) and newly generated imINPs (bottom) in type II NB lineages expressing different *UAS-nGFP* reporter constructs.

### Knockdown of Spen or Nito leads to supernumerary type II NBs and drastic increase in the expression of E(spl)mγ in type II NBs and imINPs

To investigate how the UTRs mediate post-transcriptional repression of E(spl)mγ expression, we performed an RNAi knockdown screen of RNA binding proteins with the aim of identifying proteins, knockdown of which led to increased expression of E(spl)mγ and/or supernumerary type II NB phenotypes. The supernumerary type II NB phenotype could potentially indicate increased expression of E(spl)mγ. We performed the RNAi knockdown screen in flies carrying the E(spl)mγ::GFP transgene, which not only allows for monitoring of E(spl)mγ expression in live brains but also makes flies more susceptible to supernumerary type II NB phenotypes because of an extra copy of the *E(spl)mγ* gene (Chen et al. 2021). From this screen, we identified two SPEN family RNA binding proteins, Spen and Nito, knockdown of which led to a significant increase in the number of type II NBs from 8/brain lobe in the wild type to about 114/brain lobe and 57/brain lobe, respectively. Meanwhile, the expression of E(spl)mγ in type II NBs was drastically increased by about 2-fold (Fig. 3A). Detailed examination further revealed that E(spl)mγ::GFP expression in imINPs was also significantly enhanced. In wild type, only weak E(spl)mγ::GFP is detected in about half of newly generated imINPs that are right next to the type II NB but not in any other older imINPs that are away from the NB. In Nito and Spen knockdown type II NB lineages, E(spl)mγ::GFP expression not only was dramatically increased by 4-5-fold in newly generated imINPs, but also was detected in old imINPs, with an average number of imINPs expressing E(spl)mγ::GFP increased from about 0.5/lineage in the wild type to 2-3/lineage in Spen/Nito knockdown brains (Fig. 3B).. These results indicate that Spen and Nito are required to prevent the generation of supernumerary type II NBs and to keep E(spl)mγ expression at low levels in type II NBs and imINPs.

**Figure 3.**
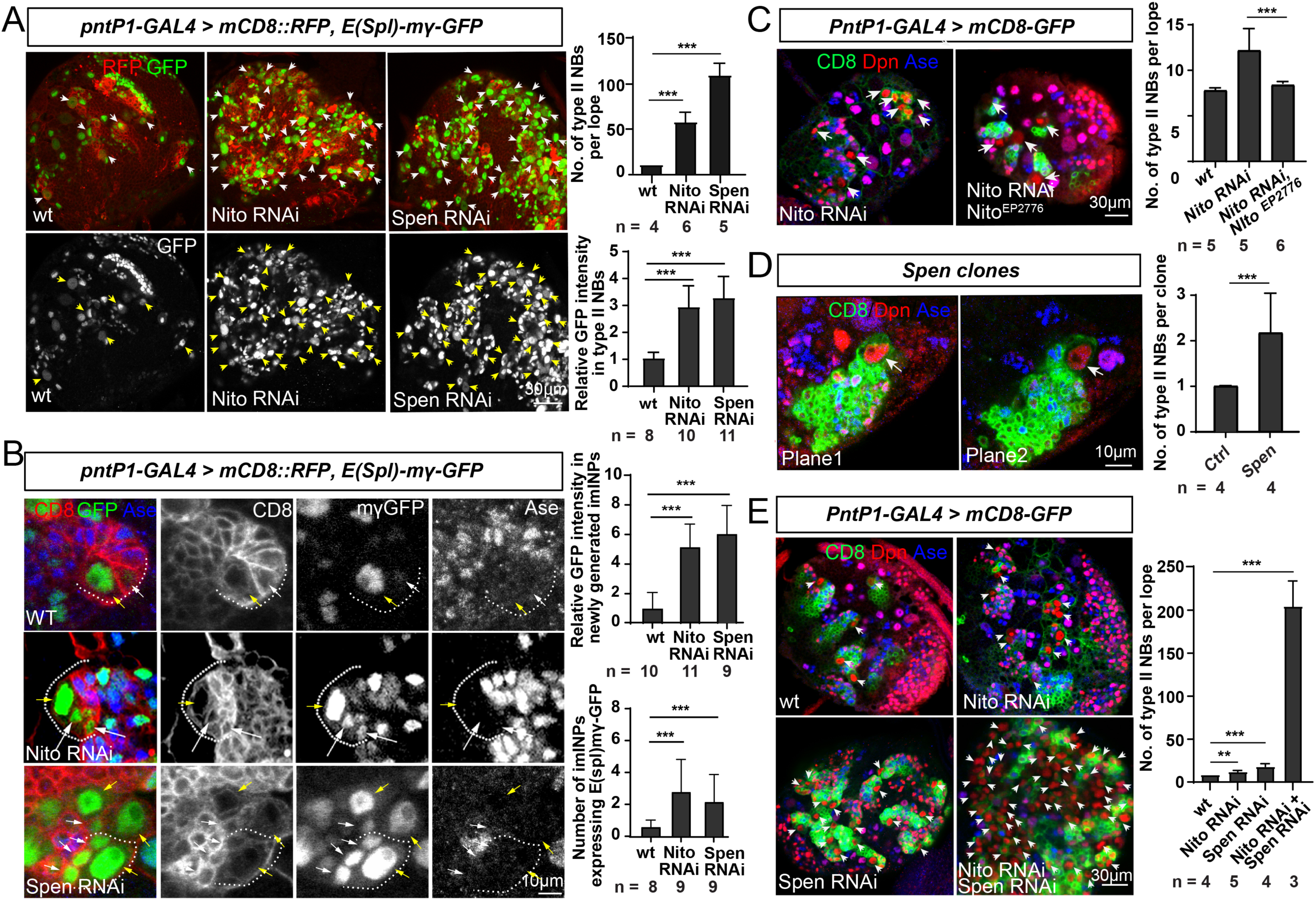
Knockdown of Spen or Nito leads to supernumerary type II NBs and increased expression of E(spl)mγ in type II NBs and imINPs. (A) Knockdown of Nito or Spen in type II NB lineages increases the number of type II NBs (arrowheads) and the expression of E(spl)mγ-GFP in type II NBs. Type II NB lineages are labeled with mCD8-RFP driven by *pntP1-GAL4*. Bar graphs to the right show the quantifications of the number of type II NBs and E(spl)mγ-GFP expression levels in type II NBs. ***, *p*<0.001, student *t*-test. (B) Nito or Spen knockdown results in persistent expression of E(spl)mγ-GFP in multiple Ase^-^ imINPs (arrows) in individual type II NB lineages (middle and bottom panels), whereas in the wild type E(spl)mγ-GFP is only detected in some (about 40%) the newly generated imINPs (arrows), but not in any older Ase^-^ imINPs (arrowheads) (top panels). Quantifications of the expression levels of E(spl)mγ-GFP in imINPs and percentages of imINPs expressing E(spl)mγ-GFP are shown in the bar graph at the bottom. ***, *p*<0.001, student *t*-test. (D) Expression of Nito from Nito^EP2776^ driven by *pntP1-GAL4* rescues the supernumerary type II NB phenotype resulting from Nito knockdown. Type II NB lineages are labeled with mCD8-GFP driven by *pntP1-GAL4*. Quantifications of the number of type II NBs are shown in the bar graph to the right. ***, *p*<0.001, student *t*-test. (E) A *spen^5^*mutant clone (labeled with mCD8-GFP) contains multiple type II NBs (arrows). Two different focal planes are shown from the same clone. Quantifications of the number of type II NBs in wild type and *spen^5^*mutant clones are shown in the bar graph to the right. ***, *p*<0.001, student *t*-test. (E) Double knockdown of Spen and Nito leads to synergistic enhancement of the supernumerary type II NB phenotypes, compared to Nito or Spen knockdown alone, in the absence of E(spl)mγ-GFP. Type II NB lineages are labeled with mCD8-GFP driven by *pntP1-GAL4*. Quantifications of the number of type II NBs in the brains with indicated genotypes are shown to the right. **, *p*<0.01 ***, *p*<0.001, student *t*-test.

To confirm that the knockdown phenotypes are not off-target effects of the *UAS-RNAi* transgenes, we examined if RNAi knockdown reduced/abolished endogenous protein expression and if the RNAi phenotypes can be rescued by expressing wild type genes or reproduced in mutant clones. Our immunostaining results showed that endogenous Nito proteins were expressed in all NBs and their progeny in wild type larval brains. Expression of *UAS-nito RNAi* driven by *pntP1-GAL4* abolished Nito expression in type II NB lineages (Fig. S2). The Nito knockdown phenotype could be largely rescued by expressing wild type Nito using an EP transgenic line *nito^EP2776^* (Fig. 3C). Additionally, *spen^5^* mutant clones also generated extra type II NBs with an average of two NBs per clone, compared to only one NB per clone in the wild type (Fig. 3D). There results confirm that the RNAi knockdown phenotypes are not off-target effects.

Spen and Nito can act antagonistically or additively/synergistically depending on the context (Chen and Rebay 2000; Kuang et al. 2000; Doroquez et al. 2007; Ma et al. 2007; Chang et al. 2008). Similar knockdown phenotypes of Spen and Nito indicate that they may function additively or synergistically. Therefore, we examined if double knockdown of Spen and Nito would lead to synergistically enhanced phenotypes. To this end, we performed the knockdown in the absence of the E(spl)mγ::GFP transgene because knockdown of Spen in the presence of the E(spl)mγ::GFP transgene already gave relatively strong phenotypes, which may make it difficult to see synergistic enhancement of the phenotypes. Knockdown of Nito or Spen alone in the absence of E(spl)mγ::GFP led to much weaker supernumerary type II NB phenotypes, with only about 12 or 15 type II NBs/brain lobe, respectively. However, double knockdown of Spen and Nito drastically increased the number of type II NBs to about 201/per brain lobe (Fig. 3E). The synergistic enhancement of the phenotypes suggests that Spen and Nito may function partial redundantly to prevent the generation of supernumerary type II NBs. The enhancement of Spen/Nito knockdown phenotypes by the E(spl)mγ::GFP transgene also indicates that Spen and Nito function in the same pathway as E(spl)mγ, which is in line with the enhanced expression of E(spl)mγ::GFP expression in Spen/Nito knockdown type II NB lineages.

### The supernumerary type II NBs resulting from Spen/Nito knockdown are derived from dedifferentiation of imINPs

The generation of supernumerary type II NBs could be due to defects in asymmetric division of type II NBs or dedifferentiation of imINPs. To rule out that the generation of supernumerary type II NBs was due to defects in asymmetric division of type II NBs, we examined the distribution of the polarity proteins Miranda (Mira) and aPKC at metaphase and anaphase of dividing Spen/Nito RNAi type II NBs. Our results show that both Mira and aPKC were segregated normally to the basal and apical cortex, respectively, at metaphase and anaphase in both Spen and Nito knockdown NBs (Figure S3), indicating that the supernumerary type II NB phenotype is not due to asymmetric division defects of type II NBs, but more likely dedifferentiation of imINPs.

Preventing dedifferentiation of imINPs requires activation of Erm, which in turn inhibits PntP1 expression in imINPs (Weng et al. 2010; Koe et al. 2014)(Janssens et al. 2017; Li et al. 2017). To determine if the supernumerary type II NBs resulting from Spen/Nito knockdown is due to dedifferentiation of imINPs, we thus examined the expression of Erm and PntP1 in imINPs. We found that knockdown of Spen or Nito both led to a significant reduction in Erm expression in imINP (Fig. 4A-B). Accordingly, the number of PntP1-positive imINPs increased significantly from 2-3/lineage in the wild type to 7-13/lineage in Spen/Nito knockdown brains (Fig. 4A, C), suggesting that the reduced expression of Erm could no longer inhibit PntP1 expression efficiently in imINPs. These data support that the supernumerary type II NBs are generated from dedifferentiation of imINPs.

**Figure 4.**
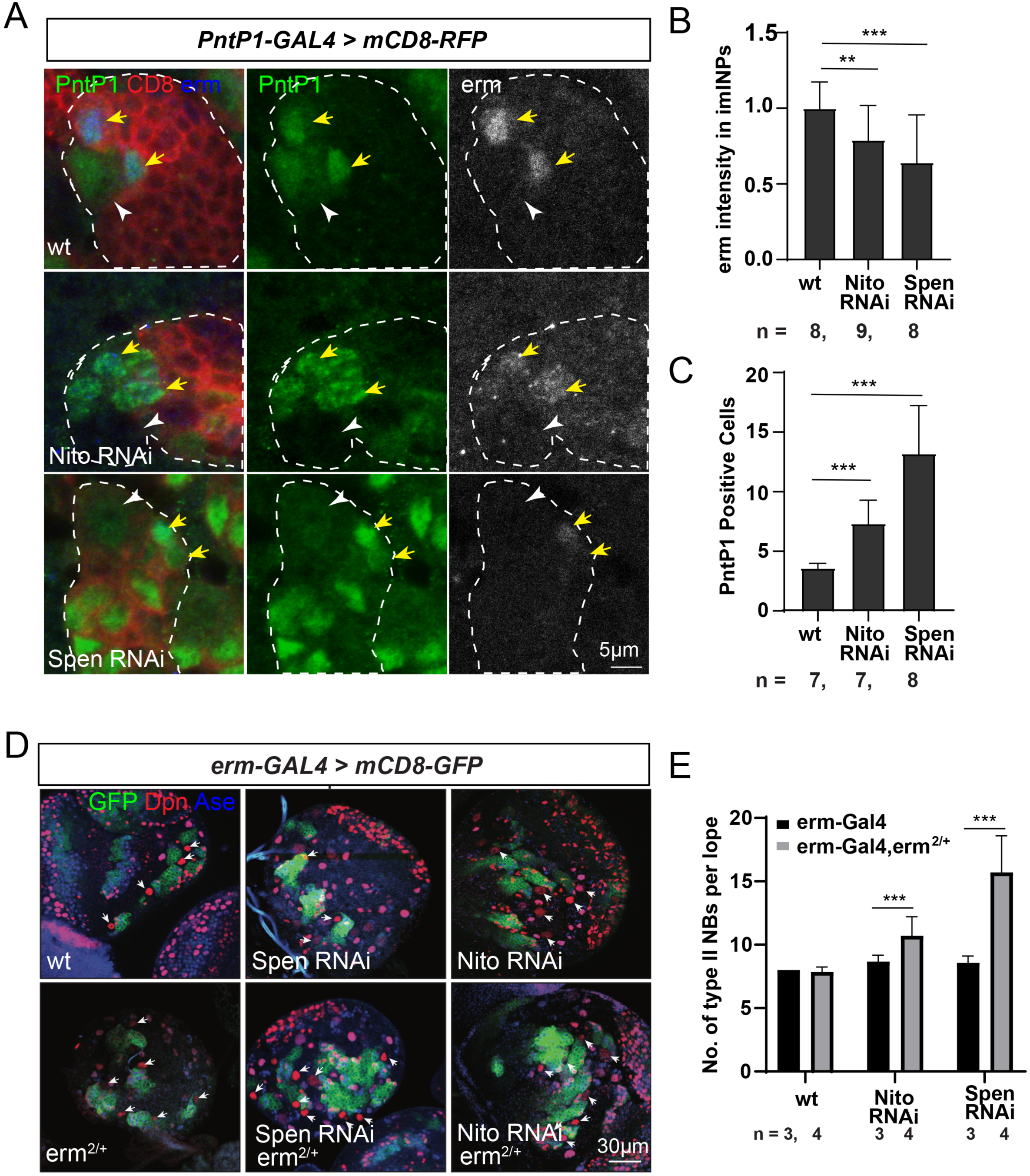
The supernumerary type II NBs resulting from Spen/Nito RNAi likely result from dedifferentiation of imINPs. (A) Knockdown of Spen or Nito leads to reduction of Erm expression in imINPs and increase in the number of PntP1-expressing imINPs. In all images, type II NB lineages are labeled with mCDE8-GFP driven by *pntP1-GAL4* and brains are counterstained with anti-PntP1 and anti-Erm antibodies. White arrowheads point to type II NBs and yellow arrows to imINPs. (B-C) Quantifications relative Erm expression levels and the number of PntP1+ imINPs in type II NB lineages with indicated genotypes. **, *p*<0.01, ***, *p*<0.001, student *t*-test. (D) The supernumerary type II NB phenotypes resulting from Spen or Nito knockdown with *erm-GAL4* is significantly enhanced in the *erm^2^/+* heterozygous mutant background. In all images, type II NB lineages are labelled with mCD8-RFP driven by *erm-GAL4* and brains are counterstained with anti-Dpn and anti-Ase antibodies. (E) Quantifications of the number of type II NBs in brains with Spen or Nito knockdown with *erm-GAL4* in the wild type or *erm^2^/+* background. ***, *p*<0.001, student *t*-test.

To more directly demonstrate that the supernumerary type II NBs were generated from dedifferentiation of imINPs, it would be ideal to knockdown Spen and/or Nito specifically in imINPs. Previously, it has been shown that the *erm-GAL4* driver on the 2^nd^ chromosome is specifically expressed in imINPs (Janssens et al. 2014). However, we recently found that this driver was also expressed in type II NBs at early larval stages in addition to imINPs (Fig. S4A). Although we were able to show that knockdown of Spen or Nito alone or together with *erm-GAL4* led to generation of supernumerary type II NBs, especially in the presence of E(spl)mγ::GFP (Fig. S4B-D), we think that these supernumerary type II NBs were more likely generated from knockdown of Spen/Nito in type II NBs instead of imINPs because Spen/Nito translationally regulates E(spl)mγ expression but E(spl)mγ mRNAs are barely detectable in imINPs (see below).Thus, the lack of a GAL4 driver specifically expressed in imINPs made it difficult to knock down Spen/Nito specifically in imINPs. Therefore, to provide additional evidence to support that Spen and Nito prevent dedifferentiation of imINPs by regulating Erm expression, we compared the phenotypes of Spen/Nito knockdown with *erm-GAL4* in the wild type and *erm^2^/+* heterozygous mutant background. Our results showed that knockdown of Nito or Spen with *erm-GAL4* in the wild type background barely led to generation of any supernumerary type II NBs. However, in the *erm^2^/+* background, knockdown of Nito or Spen with *erm-GAL4* increased the number of type II NBs to about 11 or 15 per brain lobe, respectively (Fig.4D-E). The enhancement of the phenotypes in the *erm^2^/+* background provides positive genetic interaction data to further support that dedifferentiation of imINP induced by Spen/Nito knockdown results from the reduction in Erm expression.

### The increased expression of E(spl)mγ is responsible for the generation of supernumerary type II NBs resulting from Spen/Nito knockdown

Our previous studies and others have shown that overexpression of E(Spl)mγ inhibits Erm expression and subsequently leads to dedifferentiation of imINPs (Zacharioudaki et al. 2012; Li et al. 2016; Janssens et al. 2017; Li et al. 2017). Next, we wanted to determine if the increased E(Spl)mγ expression is responsible for the generation of supernumerary type II NBs resulting from Spen/Nito knockdown. To this end, we examined if knocking down E(spl)mγ would rescue the supernumerary type II NB phenotype in Spen/Nito knockdown brains. Expression of *UAS-E(spl)mγ RNAi* significantly reduced the E(spl)mγ expression levels by 40-75% in both wild type and Spen or Nito knockdown type II NBs as indicated by the expression of E(spl)mγ::GFP (Fig. 5A, B, D, and Fig. S5A). Accordingly, the generation of supernumerary type II NBs resulting from Spen or Nito knockdown, no matter whether in the presence or absence of the E(spl)mγ::GFP), was dramatically reduced by simultaneous knockdown of E(spl)mγ, whereas knockdown of E(spl)mγ alone did not affect the number of type II NBs (Fig. 5A-C, Fig. S5B. These data demonstrate that the increased expression of E(spl)mγ is responsible for the generation of supernumerary type II NBs resulting from Spen/Nito knockdown. This is also in line with the enhancement of the Spen/Nito knockdown phenotypes by the presence of the E(spl)mγ::GFP transgene.

**Figure 5.**
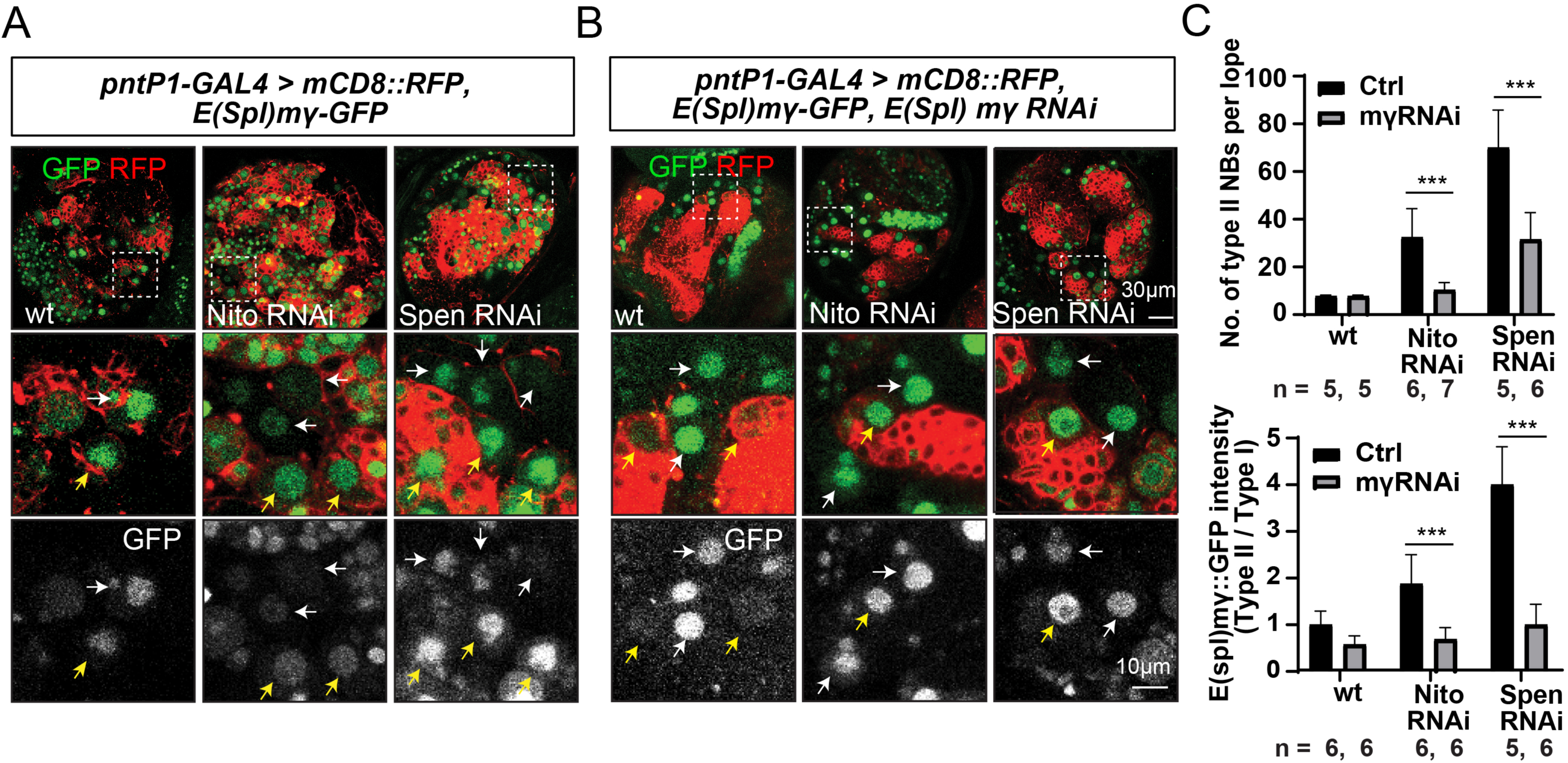
Knocking down E(spl)mγ rescues the supernumerary type II NB phenotypes resulting from Nito/Spen knockdown. In all images, type II NB lineages are labeled with mCD8-RFP driven by *pntP1-GAL4*. (A) Knockdown of Spen or Nito leads to dramatic enhancement of E(spl)mγ-GFP expression in type II NBs (yellow arrows) and supernumerary type II NBs, compared to that in wild type brains. (B) Expression of *UAS-E(spl)mγ RNAi* drastically reduces the expression of E(spl)mγ-GFP in both wild type and Spen/Nito RNAi type II NBs and largely rescues the supernumerary type II NB phenotypes resulting from Spen/Nito RNAi. Yellow arrows point to type II NBs and white arrows type I NBs in (A) and (B). (C) Quantifications of the number of type II NBs and the expression levels of E(spl)mγ-GFP in type II NBs (normalized by those in type I NBs from the same brain). ***, *p*<0.001, student *t*-test.

### Spen and Nito similarly suppress E(spl)mγ expression but not Deadpan in type I NBs

Nito is not only expressed in type II NB lineages, but also in type I NBs and their progeny. Spen was also shown to be ubiquitously expressed (Kuang et al. 2000).Thus, we also examined whether Spen or Nito knockdown would increase the expression of E(spl)mγ::GFP expression in type I NBs and induce the generation of supernumerary type I NBs. We focused our analyses on type I NB lineages in the ventral nerve cord (VNC), which contains no type II NBs. Consistent with the ubiquitous expression of Nito and Spen, we found that knockdown of Nito or Spen similarly increased the expression of E(spl)mγ::GFP in type I NBs by about 1-fold (Fig. S6A). However, we did not observe an obvious increase in the number of type I NBs in the ventral nerve cord when Spen or Nito was knocked down (Fig. S6A). Given that Prospero promotes cell cycle exit and specification of GMCs (Li and Vaessin 2000; Choksi et al. 2006), we examined the expression of Prospero. We found that Prospero was expressed normally in GMCs when Spen or Nito was knocked down type I NB lineages (Fig. S6B), which probably explains why knockdown of Spen/Nito increased the expression of E(spl)mγ expression in type I NBs but did not lead to generation of supernumerary type I NBs. These results demonstrate that Spen and Nito similarly suppress E(spl)mγ expression in type I NBs, but does not cause dedifferentiation of GMCs into type I NBs.

In addition to E(Spl)mγ, Notch also activates the expression of Deadpan (Dpn) in NBs (San-Juan and Baonza 2011). Thus, we also compared the expression level of Dpn in wild type and Spen/Nito knockdown type I NBs, but found that Dpn expression was not affected by Spen or/and Nito knockdown (Fig. S6C), indicating that Spen/Nito do not ubiquitously regulate the expression of all Notch targets, but only specific targets that contain specific binding sites for Nito/Spen (see below).

### Spen and Nito post-transcriptionally repress E(spl)mγ expression

How do Spen and Nito regulate E(spl)mγ expression? Do they function transcriptionally or post-transcriptionally? The mammalian homolog of Spen, MINT/SHARP, has been shown to inhibit Notch signaling by acting as transcriptional repressors that compete with the activated Notch intracellular domain (NCID) for binding with RBP-J (Oswald et al. 2002; Kuroda et al. 2003; Oswald et al. 2005). Therefore, we tested if Spen and Nito inhibit the transcription of *E(spl)mγ*. To this end, we generated a *E(spl)mγ* transcriptional reporter *E(spl)mγ* enhancer-nGFP, which utilizes the 946bp *E(spl)mγ* promoter upstream from the transcriptional start site to directly drive the expression of nGFP. The 947bp promoter is the promoter that drives the expression of E(spl)mγ::GFP in an endogenous expression pattern (Almeida and Bray 2005). Our results showed that knockdown of Nito or Spen did not affect the expression of *E(spl)mγ* enhancer-nGFP in type II NBs (Fig. 6A), indicating that different from mammalian Spen, *Drosophila* Spen and Nito do not regulate the transcription of *E(spl)mγ*.

**Figure 6.**
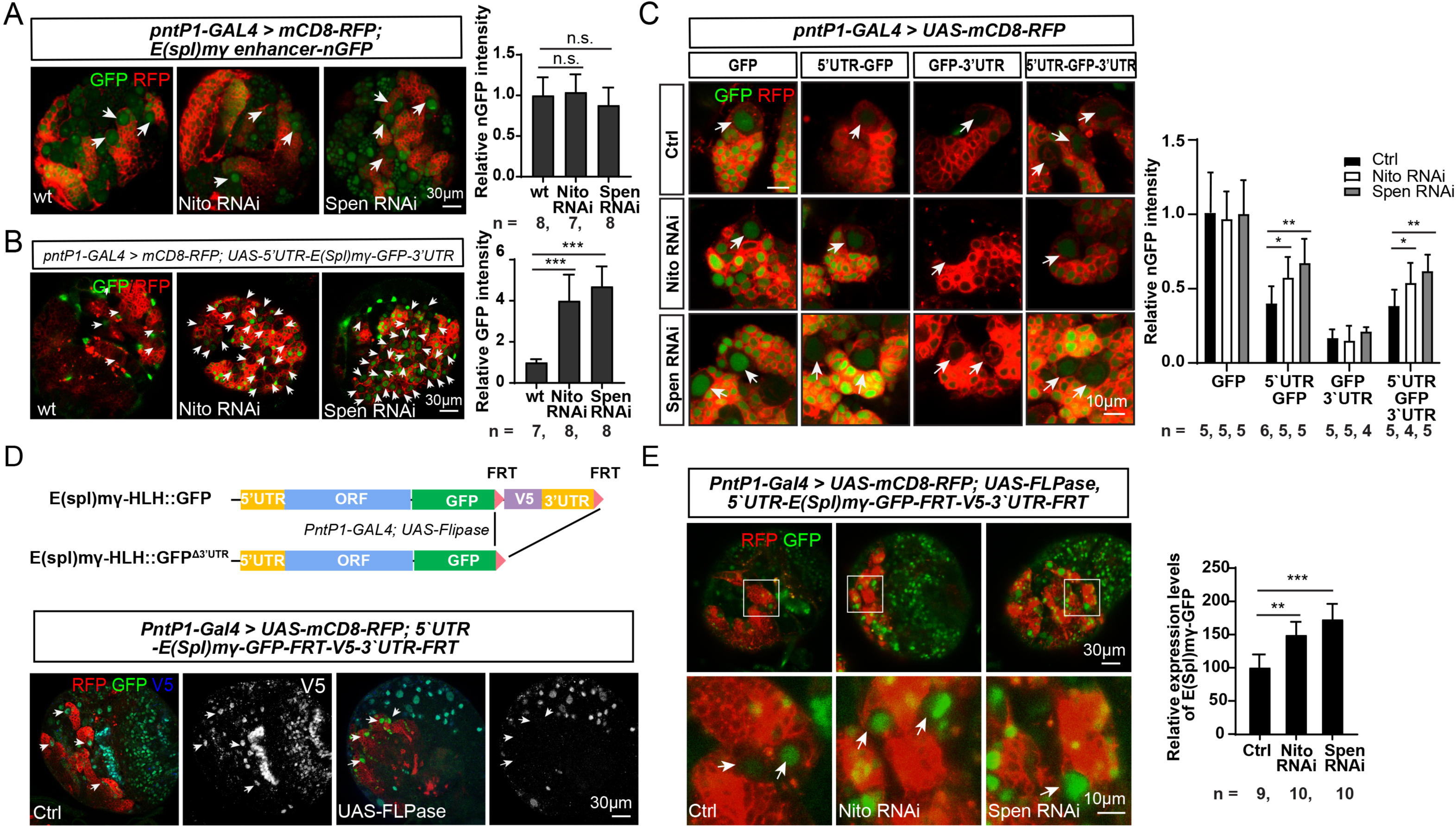
Spen/Nito post-transcriptionally regulates E(Spl)mγ expression through the 5’UTR. In all images, type II NB lineages are labeled with mCD8-RFP driven by *pntP1-GAL4*. (A) E(spl)mγ enhancer-nGFP reporter expression in type II NBs (arrows) is not affected by Nito or Spen knockdown (middle and right panels), compared to that in the wild type wild type II NBs (left panels). Quantifications of the expression levels of the nGFP reporter are shown in the bar graph to the right. n.s., not significant, student *t*-test. (B) E(spl)mγ-GFP expression from the *pUAST-5’UTR-E(Spl)mγ-GFP-3’UTR* driven by *pntP1-GAL4* is drastically enhanced by Nito or Spen RNAi (middle and right panels), compared to that in the wild type (left panels). Quantifications of the expression of E(spl)mγ-GFP are shown in the bar graph to the right. ***, *p*<0.001, student *t*-test. (C) Nito or Spen RNAi (middle and bottom rows) increases the expression of nGFP reporter from the expression of *UAS-5’UTR-nGFP* and *UAS-5’UTR-nGFP-3’UTR* but not from UAS-nGFP or *UAS-nGFP-3’UTR*, compared to the control (top row). Arrows point to type II NBs. Quantifications of nGFP expression levels are shown in the bar graph to the right. *, *p*<0.05, **, *p*<0.01, student *t*-test. (D) (Top) A schematic diagram of the E(Spl)mγ-GFP^FRT-V5-3’UTR-FRT^ construct and excision of V5 and 3’UTR from the construct by the expression of FLPase. (Bottom) The E(Spl)mγ-GFP^FRT-V5-3’UTR-FRT^ knock-in allele expresses E(Spl)mγ-GFP-V5 in type II NBs without the expression of FLPase (left panels) but loses the V5 tag expression when PntP1 drives the expression of *UAS-FLPase* (right panels). Arrows point to type II NBs. (E) Knockdown of Nito or Spen significantly increases the expression of E(Spl)mγ-GFP from the E(Spl)mγ-GFP^FRT-V5-3’UTR-FRT^ knock-in allele from which the 3’UTR together with the V5 sequence has been removed by FLPase-mediated excision of the FRT cassette (middle and right panels), compared with the control (left panels). Lower panels show enlarged views of the areas highlighted with squares in upper panels. Quantifications of the E(Spl)mγ-GFP expression levels are shown in the bar graph to the right. **, *p*<0.01, ***, *p*<0.001, student *t*-test.

To determine if Spen and Nito post-transcriptionally regulate E(spl)mγ, we then generated a *UAS-5’UTR-E(spl)mγ-GFP-3’UTR* (*UAS-E(spl)mγ^FL^-GFP*) transgenic line and used *pntP1-GAL4* to drive the expression of the transgene while knocking down Spen or Nito. In that way, the transcription of the transgene would be at the same level when Spen or Nito was knocked down. We found that, similar to the expression of E(spl)mγ-GFP under the control of the endogenous promoter, knockdown of Spen or Nito increased the expression of E(spl)mγ-GFP from the *UAS-5’UTR-E(spl)mγ-GFP-3’UTR* transgene by about 2-fold. Accordingly, Spen or Nito knockdown together with the expression of *UAS-5’UTR-E(spl)mγ-GFP-3’UTR* led to synergistic enhancement of the supernumerary type II NBs (Fig. 6B). These data indicate that Spen and Nito post-transcriptionally repress E(spl)mγ expression.

### Spen and Nito post-transcriptionally regulate E(spl)mγ expression via the 5’UTR

Next, we wanted to determine which UTRs mediate the post-transcriptional repression of E(spl)mγ by Spen/Nito. To this end, we examined how expression of the nGFP reporter from the *UAS-nGFP*, *UAS-5’UTR-nGFP*, *UAS-nGFP-3’UTR*, or *UAS-5’UTR-nGFP-3’UTR* transgenes driven by *pntp1-GAL4* would be affected by Spen or Nito knockdown. Our results show that knockdown of Spen or Nito enhanced the expression of nGFP from reporter lines containing the 5’UTR (*UAS-5’UTR-nGFP* and *UAS-5’UTR-nGFP-3’UTR*), but not from the lines without the 5’UTR (*UAS-nGFP* or *UAS-nGFP-3’UTR*) (Fig. 6C), suggesting that the 5’UTR mediates the post-transcriptional repression of E(spl)mγ expression by Spen/Nito.

To confirm that the 5’UTR mediates the post-transcriptional repression of the endogenous E(spl)mγ gene, we used the CRISPR/Cas9 technology (Jinek et al. 2012; Gratz et al. 2015) to knock in FRTs flanking the 3’UTR of the endogenous E(spl)mγ gene, which allows us to excise the 3’UTR by expressing FLPase and specifically examine if Spen/Nito knockdown would affect the expression of E(spl)mγ through the 5’UTR. Meanwhile, the GFP and the V5 coding sequences were inserted in-frame at the 3’-end of the E(spl)mγ coding sequence and between the FRT and the 5’-end of the 3’UTR, respectively. This knock-in line (5’UTR-E(spl)mγ-GFP-FRT-V5-3’UTR-FRT) expresses an E(spl)mγ-GFP-V5 fusion protein in the absence of FLPase. Upon expression of FLPase, the FLPase mediates the recombination between the FRTs and subsequent excision of the 3’UTR together with the V5 sequence, resulting in the expression of E(spl)mγ-GFP without the V5 tag (Fig. 6D). The GFP tag allows us to measure the expression level of E(spl)mγ, whereas loss of the V5 tag will indicate the excision of the 3’UTR. Our results showed that knocking down Spen or Nito still significantly increased the expression of endogenous E(spl)mγ-GFP expression by about 50-70% when the 3’UTR was excised (Fig. 6E). These results confirm that Spen and Nito post-transcriptionally repress the expression of endogenous *E(spl)mγ* gene via the 5’UTR.

### Nito directly binds to two identical novel motifs in the 5’UTR of E(spl)mγ mRNAs

To further elucidate how Spen/Nito post-transcriptionally repress E(spl)mγ expression, we examined if Spen/Nito binds to the 5’UTR. Due to the lack of appropriate reagents and the extremely large size of the Spen protein (ranging from 5458 to 5560aa depending on the isoforms), we focused the analysis on Nito. We took two distinct approaches. We first performed RNA-coimmunoprecipitation (RNA-coIP) by expressing Myc-tagged Nito in type II NB lineages using the *UAS-nito-myc* transgenic line. Using the Myc antibody, we successfully pulled down *E(spl)mγ* mRNAs (Fig. 7A), indicating that Nito interacts with *E(spl)mγ* mRNAs either directly or indirectly.

**Figure 7.**
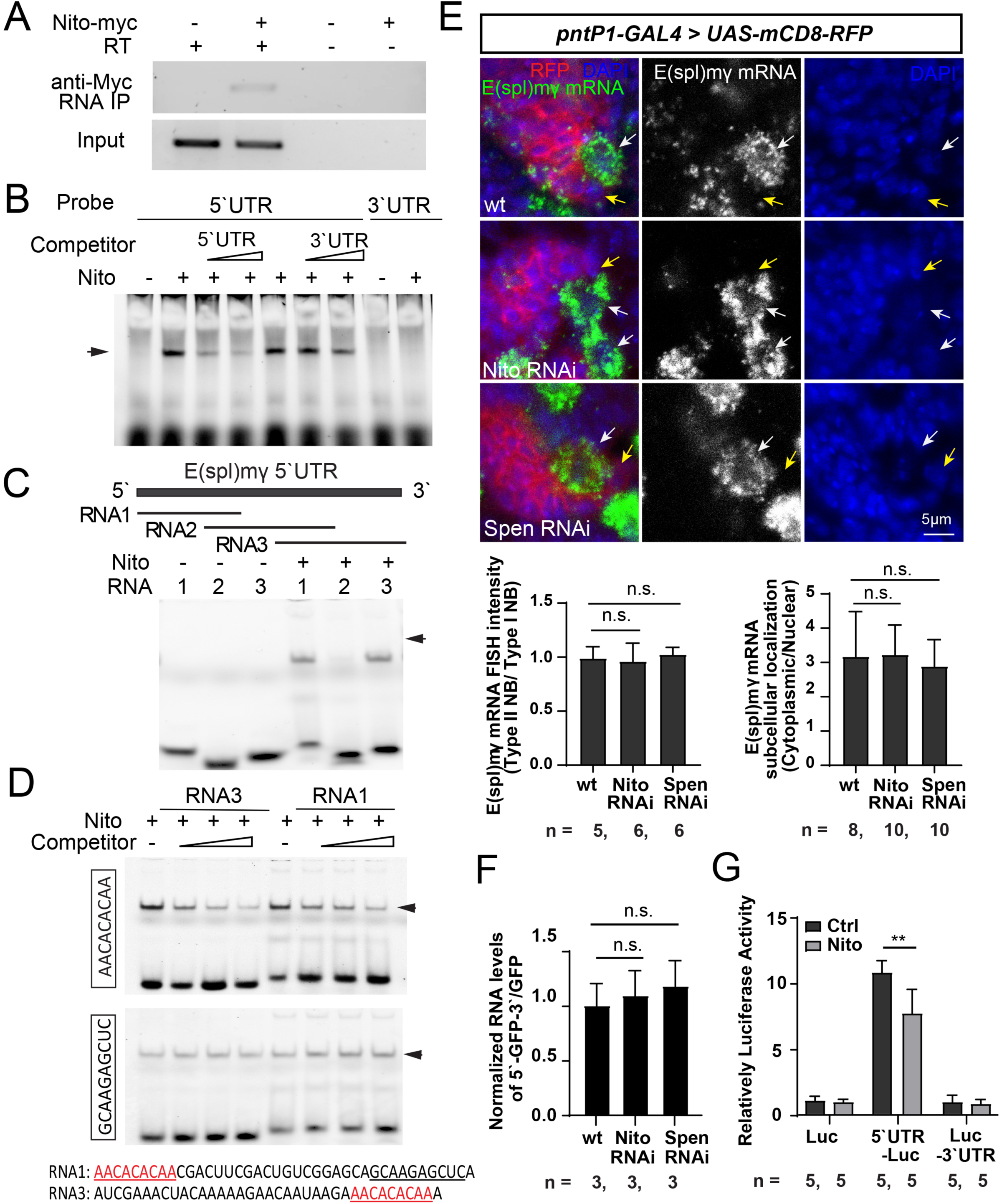
Nito regulates the translation of E(Spl)mγ transcripts by binding directly to two identical novel motifs in the 5’UTR. (A) E(Spl)mγ mRNAs are co-IPed with Nito-myc from type II NB lineages. E(Spl)mγ mRNAs were detected by RT-PCR after RNA-coIP. (B) RNA-EMSA shows Nito binds to the E(spl)mγ 5’UTR but not the 3’UTR. The binding can be competed with the cold probe of the 5’UTR but not the 3’UTR. The arrow points to retarded bands. (C) RNA-EMSA shows Nito binds to the fragment RNA1 and RNA2 but not RNA3 from the E(spl)mγ 5’UTR. The arrow points to retarded bands. (D) RNA-EMSA shows binding of Nito to the fragments RNA1 and RNA3 can be competed by a cold probe of their common sequence (in red) found in RNA1 and RNA3 but not by a cold probe of the underlined sequence from the 3’-end of RNA1. (E) In situ hybridization chain reaction results show that compared to the wild type (upper panels), Nito or Spen knockdown does not affect E(Spl)mγ mRNA levels or their distribution in the cytoplasm and nucleus in type II NBs (white arrows) or degradation of E(Spl)mγ mRNA in newly generated imINPs (yellow arrows). Type II NB lineages are labeled with mCD8-RFP driven by *pntP1-GAL4* and counter-stained with DAPI. Bar graphs at the bottom show quantifications of E(Spl)mγ mRNA in situ signal intensities (normalized by those in type I NBs) (upper) and the ratio of the signal intensities in the cytoplasm vs in the nucleus (bottom). n.s., not significant, student *t*-test. (F) qRT-PCR results show nGFP mRNA levels (normalized by eEF1a mRNA levels) from the expression of *UAS-5’UTR-nGFP-3’UTR* driven by *pntP1-G*AL4 are not affected by Spen or Nito RNAi in type II NB lineages. n.s., not significant, student *t*-test. (G) Luciferase assays show that Nito inhibits the translation of the transcripts of 5’UTR-Luc, but not the transcripts of Luc alone or Luc-3’UTR *in vitro*. **, *p*<0.01, student *t*-test.

To further determine if Nito binds to the 5’UTR directly, we then performed RNA-electrophoretic mobility shift (RNA-EMSA). Using Nito protein expressed *in vitro* and synthesized fluorescein-labeled 5’UTR, we found that Nito bound to the 5’UTR as indicated by the presence of a retarded band, but not the 3’UTR. The binding to the 5’UTR could be competed with a cold probe of 5’UTR but not by a cold probe of 3’UTR (Fig. 7B). These results demonstrate the Nito binds to the 5’UTR directly. To further map the binding motifs of Nito, we performed similar gel-mobility assays using three smaller fragments with overlapping sequences from the 5’UTR. We found that Nito could bind to the fragment from the 5’-end and the one from the 3’-end but not the one from the middle of the 5’UTR (Fig. 7C). Visual examination of the sequence of these two fragments revealed that these two fragments share a sequence 5’-AACACACAA-3’, which is located at both the 5’-end and 3’-end of the 5’UTR. Further competition assays showed that a cold RNA probe containing this sequence could compete with the full-length 5’UTR for binding with Nito, but not a probe containing an irrelevant sequence from the middle of the 5’UTR (Fig. 7D), confirming that 5’-AACACACAA-3’ is the Nito binding motif.

### Nito represses translation of E(spl)mγ mRNAs

RNA binding proteins can post-transcriptionally regulate the expression of target genes by regulating the stability, nuclear export, or translation of the transcripts (Glisovic et al. 2008; Ceci et al. 2021). To examine if Spen/Nito regulate the stability of mRNAs, we performed in situ hybridization chain reaction of *E(spl)mγ* mRNAs in Spen/Nito knockdown type II NB lineages as well as quantitative real-time RT-PCR to compare the level of nGFP mRNAs transcribed from *UAS-5’UTR-nGFP-3’UTR* driven by *pntP1-GAL4* in the wild type and Spen or Nito RNAi brains. Our results show that the overall in situ signal intensities in type II NBs (normalized by signal intensities in type I NBs in the same brain) were not significantly changed by Spen or Nito knockdown (Fig. 7E). Furthermore, like in the wild type, *E(spl)mγ* mRNA in situ signals in imINPs in Spen/Nito knockdown type II NB lineages were largely gone except for a few dots (Fig. 7E). These data indicate that Spen and Nito do not regulate the stability of *E(spl)mγ* mRNAs in type II NBs or degradation of *E(spl)mγ* mRNAs in imINPs. Consistently, real-time RT-PCR results showed that knocking down Spen or Nito did not significantly change the nGFP mRNA levels transcribed from the *UAS-5’UTR-nGFP-3’UTR* transgene (Fig. 7F), further confirming that binding of Spen or Nito to the 5’UTR do not regulate the stability of mRNAs.

To determine if Spen and Nito regulate nuclear export of *E(spl)mγ* mRNAs, we also compared the ratio of their *in situ* signals in the nucleus *v.s.* in the cytoplasm. Our results showed that Spen or Nito knockdown did not significantly change the distribution of *E(spl)mγ* mRNAs in the nucleus and the cytoplasm. In both wild type and Spen/Nito RNAi type II NBs, *E(spl)mγ* mRNAs were predominantly localized to the cytoplasm (Fig. 7E). Therefore, Spen and Nito do not regulate nuclear export of *E(spl)mγ* mRNAs either.

To examine if Nito regulates the translation of *E(spl)mγ* mRNAs, we generated luciferase reporter constructs by fusing the 5’UTR or the 3’UTR sequence with the luciferase coding sequence and performed luciferase assays. We found that adding Nito proteins to the *in vitro* cell-free translational system significantly suppressed the expression of Luciferase from the luciferase transcripts fused with the 5’UTR by about 30% but not with the 3’UTR or without the UTRs (Fig. 7G), suggesting that Nito translationally suppresses E(spl)mγ expression by binding to the 5’UTR. In addition, in the *in vitro* translational system, fusion of the 5’UTR to the luciferase coding sequence dramatically boosted the expression of Luciferase. Our *in vivo* nGFP reporter results indicate that the 5’UTR overall exerts an inhibitory effect on gene expression but also contains positive regulatory motifs. It is possible that the negative regulatory mechanisms (e.g. promoting mRNA degradation) may not be functioning in the *in vitro* translational system due to inhibition of RNAse in the system. Therefore, the positive regulatory effect becomes more obvious.

## DISCUSSION

In this study, we demonstrate that, in addition to Numb-mediated degradation of Notch in imINPs, fate commitment of INPs also requires the expression of the Notch target E(Spl)mγ in type II NBs to be kept at low levels by Spen and Nito. Spen and Nito reduce the expression of E(Spl)mγ by binding to two identical novel motifs in the 5’UTR to repress the translation of E(Spl)mγ transcripts (Fig. S7A). The low expression in type II NBs ensures that the E(Spl)mγ proteins inherited by newly generated imINPs will be rapidly removed from imINPs right after division of the NB, which in turn allows Erm to be activated by PntP1 and promote INP maturation. In the absence of Spen/Nito, de-repression of translation leads to increased expression of E(Spl)mγ in type II NBs and subsequent delayed clearance of E(Spl)mγ from imINPs and dedifferentiation of imINPs into type II NBs due to inhibition of Erm expression (Fig. S7B). This Spen and Nito-mediated translational regulation of E(spl)mγ expression in type II NBs acts in concert with Numb-mediated degradation of Notch to ensure Notch signaling in imINPs to be rapidly shut off right after the asymmetric division of type II NBs. The Numb-mediated degradation of Notch turns off the transcription of E(spl)mγ in imINPs so that no new E(spl)mγ mRNAs will be produced, whereas the Spen and Nito-mediated translational repression of E(spl)mγ expression in type II NBs ensures no excessive amount of E(spl)mγ proteins will be inherited by imINPs. Defects in either of the mechanisms will similarly lead to ectopic/prolonged expression of E(spl)mγ in imINPs and subsequent dedifferentiation of imINPs. Together, our work reveals a novel post-transcriptional mechanism that is as critical as Numb-mediated degradation of Notch in INP cell fate commitment.

In mammals, the Spen homolog, SHARP/MINT, and the Nito homolog, RBM15), have previously been shown to regulate Notch signaling, but they mainly function as transcriptional repressors that competes with the activated Notch intracellular domain (NCID) for binding with RBP-J (Oswald et al. 2002; Kuroda et al. 2003; Oswald et al. 2005). In the absence of Notch activation, SHARP/MINT binds to RBP-J and suppresses the transcription of Notch target genes. Upon activation of Notch, NCID binds to RBP-J by displacing SHARP/MINT and activates transcription of Notch target genes. Similarly, RBM15 interacts with RBP-J to regulate Notch signaling (Ma et al. 2007).In mammals, SHARP/MINT/, inhibits Notch signaling by acting as a transcriptional repressor In *Drosophila*, Spen can also antagonize or promote Notch signaling depending on the tissue types. In the developing eye, Spen antagonizes Notch signaling (Doroquez et al. 2007), whereas in hematopoietic cells, glial cells, and self-renewing intestinal stem cells, Spen promotes Notch signaling (Jin et al. 2009; Andriatsilavo et al. 2018; Girard et al. 2020). However, the underlying mechanisms are not well understood. In this study, we demonstrate that in *Drosophila* NBs, Spen and Nito function differently from their mammalian homologs to antagonize Notch signaling. First, instead of functioning as an ON/OFF switch to completely shut off Notch target expression, *Drosophila* Spen/Nito function as a dimmer switch to keep the expression of E(spl)mγ at low levels. Second, instead of functioning as transcriptional repressors, *Drosophila* Spen/Nito function post-transcriptionally. The post-transcriptional function of *Drosophila* Spen/Nito is consistent with their lack of the RBP-J interacting domain (RBP-ID) found in the mammalian homologs. There is no evidence supporting direct interactions between Spen/Nito and Su(H) (the *Drosophila* RBP-J) in *Drosophila* (Maier 2019). Instead, Hairless is considered the functional equivalent of SHARP/MINT and interacts with Su(H) (*Drosophila* homolog of RBP-J) through its NT box to repress the transcription of Notch targets in *Drosophila* (Morel et al. 2001; Barolo et al. 2002). Therefore, although *Drosophila* Spen and Nito share the RRM and SPOC domains with their mammalian homologs, their molecular mechanisms in regulating Notch signaling are different. In fact, post-transcriptional functions have never been reported for Spen/SHARP/MINT except that mouse MINT regulates stability of Xist mRNAs (Robert-Finestra et al. 2021; Rodermund et al. 2021). However, it is not clear if MINT regulates Xist mRNA stability directly or indirectly.

Our results show that Spen and Nito post-transcriptionally downregulate E(spl)mγ expression through the 5’UTR. 5’ UTRs harbor various regulatory elements that can bind to RNA-binding proteins, microRNAs, or ribosome entry sites (IRESs), upstream open reading frames (uORFs), and secondary structures (Ryczek et al. 2023; Razumova et al. 2025). These elements and structural components regulate translation initiation, mRNA stability, and pre-mRNA splicing. *Drosophila* Nito and Spen have been shown to regulate splicing of *sxl* pre-mRNAs (Lence et al. 2016; Kan et al. 2017; Knuckles et al. 2018), as well as localization of Xist RNAs (Bowness et al. 2022). In mammals, SHARP/MINT and RBM15 also regulates mRNA splicing, stability, nuclear export through association with messenger ribonucleoprotein (mRNP) complexes or nuclear export receptors (Hiriart et al. 2005; Lindtner et al. 2006; Uranishi et al. 2009; Zhang et al. 2015; Rodermund et al. 2021). Since *E(spl)mγ* transcripts do not contain introns, Spen or Nito cannot be involved in regulating splicing of *E(spl)mγ* pre-mRNAs. Our in situ and qRT-PCR results did not support involvement of Spen and Nito in regulating mRNA stability or nuclear export either. Instead, our *in vitro* translational assays using Luciferase as a reporter show that Nito represses translation by binding to the 5’UTR. Although we do not have direct evidence to demonstrate Spen translationally represses E(spl)mγ expression due to the large size of Spen proteins and a lack of proper reagents, our data that knockdown of Spen enhanced the expression of nGFP from *UAS-5’UTR-nGFP* or *UAS-5’UTR-nGFP-3’UTR* transgenes but did not affect the stability and nuclear export of nGFP and endogenous E(spl)mγ mRNAs also strongly suggests that Spen translationally regulates E(spl)mγ expression. Interestingly, compared to the increase of E(Spl)mγ expression in Spen/Nito knockdown type II NBs, the suppression of the Luciferase reporter expression *in vitro* by Nito is much milder. The stronger enhancement of E(Spl)mγ expression in Spen/Nito knockdown type II NBs could be that small increase in the translation of E(Spl)mγ transcripts resulting from Spen/Nito knockdown could be amplified by the positive feedback loop between E(Spl)mγ and the super-elongation complex as demonstrated before (Liu et al. 2017). SPEN family proteins often bind to RNAs at specific RNA structures like stem loops, preferentially within the 3’UTR (Arieti et al. 2014) rather than defined sequences. However, in some cases, Spen proteins can also bind at specific sequences of some target RNAs. For example, Spen/SHARP/MINT binds to Xist and endogenous retroviral (ERV) elements at A-repeats (Chu et al. 2015; Carter et al. 2020) and to the long non-coding RNA SRA (steroid receptor activator) at the H12 to H13 sequence. In this study, we found that Nito binds to two identical novel motifs containing the sequence 5’-AACACACAA-3’ at the 5’UTR E(Spl)mγ mRNAs. This sequence likely determines the specificity of target RNAs of Nito and Spen. In support of this notion, our results show that Spen and Nito do not regulate the expression of another Notch target Dpn that does not contain this sequence in either its 5’UTR or 3’UTR. Interestingly, the 3’UTR of E(Spl)mγ mRNAs also contains a single copy of this sequence. Yet, the 3’UTR neither bound to Nito in gel shift mobility assays nor mediated the suppression of nGFP reporter expression by Spen or Nito. A recent study reports that high-affinity binding of Spen to Xist requires at least four segments of A-repeats (Button et al. 2024). It is possible that efficient binding of Nito to E(Spl)mγ mRNAs also requires at least two copies of the motif. However, the additional copy of the motif in the 3’UTR does not seem to contribute to the suppression of E(Spl)mγ expression by Nito and Spen as the nGFP reporter expression from the *UAS-5’UTR-nGFP-3’UTR* transgene didn’t exhibit stronger inhibition by Spen or Nito than from the *UAS-5’UTR-nGFP* transgene. Exactly how Spen and Nito regulate translation of *E(spl)mγ* mRNAs by binding to these novel motifs remains to be investigated in the future. It doesn’t appear this novel motif will form stem loops or other secondary structures. One interesting idea to test could be that binding of Nito/Spen to both copies of the motifs in the 5’UTR may promote the 5’UTR to form a loop, which may limit the access of the ribosome to the start codon or assembly for translation initiation complex.

Recent studies have shown that Nito/RBM15 is a subunit of the N^6^-methyladenosine (m^6^A) writer complex, which consists of a catalytic subunit, a heterodimer of METTL3 and METTL14 and a regulatory subunit that includes WTAP, VIRMA, ZC3H13, HAKAI, and RBM15 (Patil et al. 2016; Knuckles et al. 2018; Su et al. 2022). m^6^A is the most abundant internal RNA modification in eukaryotic mRNAs, with thousands of human and mouse genes modified by m^6^A, including several Notch effectors (Dominissini et al. 2012; Meyer et al. 2012). m^6^A modifications can regulate splicing, localization, translation, and degradation of mRNAs (Zaccara et al. 2019; Murakami and Jaffrey 2022). Surprising, our unpublished results show that loss of the components of the m^6^A modification pathway did not lead to supernumerary type II NB phenotypes as the loss of Spen or Nito did. Considering that m^6^A modifications often occur in the 3’UTRs at sites close to the stop codon (Meyer et al. 2012), whereas Spen/Nito function through the 5’UTR of *E(spl)mγ* transcripts, it probably makes sense that m^6^A modifications are not involved in translational regulation of *E(spl)mγ* expression by Spen/Nito. Therefore, Spen and Nito likely regulate translation of *E(spl)mγ* mRNAs through other m^6^A modification-independent mechanisms.

In addition to mediating translational repression by Spen and Nito, the 5’UTRs likely also mediate other regulatory mechanisms. Our results show that the *UAS-5’UTR-nGFP-3’UTR* transgene expresses higher nGFP levels than *UAS-nGFP-3’UTR*, indicating that the 5’UTR may also contain elements mediating positive regulation of E(spl)mγ expression, which is consistent with our in vitro translational assay results showing that including the 5’UTR dramatically enhances the expression of Luciferase reporter expression. Furthermore, the nGFP reporter transgene containing the 3’UTR alone also expressed much lower levels of nGFP than the transgene containing the nGFP coding sequence alone, indicating that the 3’UTR is also critical for keeping the expression of E(spl)mγ low. Previous studies with GFP or lacZ reporter assays showed that the 3’-UTRs of *E(spl)* genes harbor binding sites for multiple classes of miRNAs and mediate the suppression by miRNAs (Lai and Posakony 1997; Lai et al. 1998; Lai et al. 2005). Thus, post-transcriptional regulation of E(spl)mγ expression may involve multiple different mechanisms, which likely function together to tightly control the expression of E(spl)mγ. Further investigation into these mechanisms will be essential to fully understand how the expression of E(spl)mγ is controlled post-transcriptionally.

In summary, our work reveals that in addition to Numb-mediated degradation of Notch, Spen/Nito-mediated translational control of the expression of Notch target E(Spl)mγ is equally critical for terminating Notch signaling in imINPs in a timely manner and proper cell fate specification of INPs. Our work also elucidates a novel post-transcriptional mechanism of the regulation of Notch target expression by Spen/Nito. Given that Spen and Nito function as tumor suppressors in many different cancers, including glioblastoma, findings from this work could also provide novel insights into the development of various cancers related with Spen/Nito.

## MATERIALS AND METHODS

### Fly husbandry

*UAS-Nito RNAi* (y^1^ v^1^; P{TRiP.HMJ02081}attP40, Bloomington Drosophila Stock Center (BDSC), #56852), *UAS-Spen RNAi* (y^1^ sc* v^1^ sev^21^; P{TRiP.HMS00276}attP2, BDSC #33398), and *UAS-E(Spl)m*γ *RNAi* (y^1^ v^1^; P{TRiP.JF02000}attP2, BDSC #25978), were used for RNAi knockdown of their respective genes; Spen5, FRT40A/CyO, P{lacZ.w+}276 (BDSC #8734) and *hs-FLPase, elav-GAL4, UAS-mCD8-GFP; tub-GAL80, FRT40A* (derived from a combination of BDSC #5146 (*P{w[+mW.hs]=GawB}elav[C155], P{w[+mC]=UAS-mCD8::GFP.L}Ptp4E[LL4], P{ry[+t7.2]=hsFLP}1*)and BDSC #5192 (*y[1] w[*]; P{w[+mC]=tubP-GAL80}LL10 P{ry[+t7.2]=neoFRT}40A/CyO*) for generating *spen* mutant clones; *Nito^EP2776^* (BDSC #27009) for rescuing Nito RNAi phenotypes; *UAS-nito-myc/CyO* (BDSC #94916) for RNA co-immunoprecipitation; *erm^2^/cyo* (Weng et al. 2010) for knocking down Spen/Nito in *erm* heterozygous mutant background; *UAS-mCD8-GFP* and *UAS-mCD8-RFP* for labeling NB lineages; *UAS-brat RNAi* y[1] sc[*] v[1] sev[21]; P{y[+t7.7] v[+t1.8]=TRiP.HMS01121}attP2, BDSC #34646) for knocking down Brat and boosting the population size of type II NBs for co-immunoprecipitation of mRNAs bound to Nito; E(Spl)mγ-GFP (Almeida and Bray 2005) as a reporter for the expression of E(Spl)mγ expression and for adding an additional copy of the E(Spl)mγ gene; *pntP1-GAL4* (Zhu et al. 2011), *erm-GAL4 (II)* (Pfeiffer et al. 2008; Janssens et al. 2014), and *insc-GAL4* (Luo et al. 1994) for driving UAS-transgenes in type II NB lineages, imINPs, and all NBs, respectively. Flies were fed with standardized food and raised at 25°C with except for RNAi knockdown, which were performed at 29°C to enhance the efficiency.

### Generation of transgenic lines

Construction of plasmids was described in detail in Supplementary information. *pUASTattB-E(Spl)m*γ*^ORF^*, *pUASTattB-E(Spl)m*γ*^FL^*, *pUAST-5’UTR-E(Spl)m*γ*-GFP-3’UTR*, *pH-Stinger-E(spl)m*γ *promoter* transgenic lines were generated by injecting plasmids into *w^1118^* preblastoderm embryos for P-element-mediated random integration into the fly genome.

*pUASTattB-5’UTR-nGFP-3’UTR*, *pUASTattB-5’UTR-nGFP, pUASTattB-nGFP-3’UTR, and pUASTattB-nGFP* transgenic lines were generated by injecting plasmids into the embryos of the fly line, y^1^ M{vas-int.Dm}ZH-2A w*; PBac{y^+^-attP-9A}VK00020 (BDSC # 24867) for integration into the same attP-9A site. Injections of plasmids into fly embryos were performed by Rainbow Transgenic Flies, Inc. (Camarillo, CA, USA).

### Generation of the E(Spl)mγORF-eGFP-FRT-V5-3’UTR-FRT knock-in line

Construction of guide RNA (gRNA) expression plasmids and the pHD-5’Arm-E(Spl)mγORF-eGFP-FRT-V5-3’UTR-FRT-DsRed-3’Arm donor vector was described in detail in the Star Methods. The donor vector together with the gRNA expression plasmids were injected into preblastoderm embryos of the Vas-Cas9 line (w[1118]; PBac{y[+mDint2] GFP[E.3xP3]=vas-Cas9}VK00037/CyO, P{w[+mC]=Tb[1]}Cpr[CyO-A], BDSC #56552). The injected adult flies were outcrossed with *w^1118^* flies. Progeny were first screened for dsRed+ eyes, followed by PCR genotyping, for identifying the knock-in lines.

### RNA Co-Immunoprecipitation

Co-immunoprecipitation of E(Spl)mγ mRNAs and Nito-Myc was performed as previously described (Connell et al. 2024). Third instar larval brains expressing *UAS-nito-myc* and *UAS-brat RNAi* driven by *pntP1-Gal4* were dissected in the RNase-free PBS and lysed in RNase-free cell lysis buffer (10 mM HEPES, pH7.0, 100 mM KCl, 5, mM MgCl2, 0.25% NP40, 0.25% Triton X-100, 0.01% SDS, with 1x Protease Inhibitor Cocktail (Catalog #78437, ThermoFisher Scientific) and 100U/ml Ribolock RNase Inhibitor (Catalog #EO0382, ThermoFisher Scientific). The protein concentrations of the lysates were quantified using a Pierce BCA Protein Assay Kit (Catalog #23227, ThermoFisher Scientific). Cell lysates were diluted in the IP buffer (50 mM Tris.HCl, pH7.4; 150 mM NaCl, 1 mM MgCl2, 0.05% NP-40, 1 mM DTT with 100U/ml Ribolock RNase inhibitor and 1x proteinase inhibitor cocktail) and incubated with Pierce Anti-c-Myc Magnetic Beads (Catalog #88842, ThermoFisher Scientific) at 4°C overnight. After washing with the IP buffer for five times, the total RNAs from the complexes were isolated using RNAzol RT (Catalog #RN 190, Molecular Research Center, Inc., Cincinnati, OH) and then used for generating a cDNA library with MMLV H-Reverse transcriptase (Catalog #M1701, Promega, Madison, WI) and Random primers (Catalog # 48190011, Invitrogen™) after DNase I (Catalog # M0303L, New England Biolabs, Ipswich, MA) treatment. A negative control without MMLV H-reverse transcriptase was prepared for each sample. PCRs were performed using DreamTaq Hot Start Green PCR Master Mix (Catalog #K9022, ThermoFisher Scientific) with the primers ORF_Fw and ORF_Rv. The RNA co-IP experiment was repeated twice to ensure reproducibility of the results.

### In Situ Hybridization Chain Reaction (HCR)

In situ HCR (Dirks and Pierce 2004; Tsuneoka and Funato 2020) was performed using the HCR™ Gold RNA-FISH kit (Molecular Instruments, Los Angeles, California) according to manufacturer’s instructions. Briefly, third instar larval brains were dissected in PBS and fixed in 100 mM PIPES (pH 6.9), 1 mM EGTA, 0.3% Triton X-100 and 1 mM MgSO4 containing 4% formaldehyde for 23 minutes. Fixed samples were washed with PBST containing 0.3% Triton X-100. Samples were pre-hybridized with hybridization buffer at 37°C for 1 hour and then incubated with 10 nM E(spl)mγ HCR probe sets (Molecular Instruments) at 37°C overnight. After hybridization, samples were washed 2 x 15 min with washing buffer and 3 x 5 min with 5 x SSCT (5 x SSC, 0.1% Tween 20). Samples were equilibrated in amplification buffer at 25°C for 30 min and then incubated with amplification buffer with 10pmol snap-cooled H1 hairpins and 10pmol snap-cooled H2 in the dark at 25°C for overnight. Samples were washed 4 x 15 min with SSCT next day. If antibody staining was required, samples were re-fixed in 100 mM PIPES (pH 6.9), 1 mM EGTA, 0.3% Triton X-100 and 1 mM MgSO4 containing 4% formaldehyde for 15 min before immunofluorescent staining procedures as described below. The E(spl)mγ mRNA in situ signal intensity in Type II neuroblasts were quantified using Zen Software (Zeiss).

### RNA electrophoretic mobility shift (RNA-EMSA) assays

Nito protein was expressed *in vitro* from the template plasmid pOT2-T7-Nito with the T7 promoter and the full-length Nito cDNA [Catlog #GH11110, *Drosophila* Genomic Resource Center (DGRC)] using the TNT T7-Quick Coupled Transcription/Translation System (Catalog #L1170, Promega) according to the manual. The fluorescein labeled E(spl)mγ 5’UTR and 3’UTR RNAs were synthesized *in vitro* by oligonucleotide-directed transcription using the T7 RNA Synthesis Kit (Catalog #E2040, New England Biolabs) and Fluorescein RNA Labeling Mix (Catalog #11685619910, Sigma-Roche, St. Louis, MO) according to the manual. The T7-5′UTR and T7-3′UTR templates for *in vitro* transcription were generated by PCR using the primers T7-5’UTR_Fw and T7-5’UTR_Rv, and T7-3’UTR_Fw and T7-3’UTR_Rv, respectively. Sequences of the primers can be found in Supplementary Table S1.

The binding reaction were performed by incubating the fluorescein labeled UTR RNAs with 5ul TNT-T7 Nito expressing reaction in the RNA-EMSA binding buffer supplemented with 10mM HEPES (PH 7.3), 20mM KCl, 1mM MgCl2, 1mM DTT, 10% Glycerol for 30 min at room temperature. The RNA-protein complex was resolved on a 5% non-denaturing polyacrylamide Mini-PROTEAN TBE Precase Gel (Catalog #4565013, Bio-Rad Laboratories, Hercules, CA) in 0.5x TBE. The gels were scanned and analyzed with ChemiDoc XRS (Bio-Rad Laboratories).

### Luciferase assays

For testing if Nito inhibits the translation of E(Spl)mγ mRNAs, Nito protein was expressed *in vitro* using the TNT T7-Quick Coupled Transcription/Translation System (Catalog #L1170, Promega, Madison, WI) according to the manual. Meanwhile, the 5′UTR and 3′UTR of E(Spl)mγ were cloned into the pT7-Luciferase vector to generate *pT7-5′UTR-Luciferase* and *pT7-Luciferase-3′UTR* plasmids (see Supplementary information for detailed description of the construction of the plasmids), which were used for *in vitro* transcription of 5′UTR-Luciferase and Luciferase-3′UTR mRNAs using the T7 RNA Synthesis Kit (Catalog # E2040, New England Biolabs). An equal amount of 5′UTR-Luciferase or Luciferase-3′UTR mRNAs mRNAs were added to the rabbit reticulocyte lysate (Catalog #L4960, Promega) with or without the TNT-T7 expressed Nito protein for *in vitro* translation at 30°C for 90min. The luciferase activities were measured with BioTek Synergy Neo2 Hybrid Multimode Reader (Agilent, Santa Carla, CA) using the ONE-Glo™ EX Luciferase Assay System (Catalog #E8130, Promega) according to the instruction of the manufacture.

### Immunofluorescent and confocal microscopy

Larval brains were dissected in PBS and fixed in 4% of formaldehyde in PBS for 20min at room temperature. After washing in PBT (PBS with 0.3% of Triton-X100), larval brains were incubated with the primary antibodies in PBT with 5% normal goat serum at 4°C overnight, followed by washing with PBT. The brains were then incubated with the fluorophore-conjugated secondary antibodies in PBT at room temperature for 2 hours. The stained brains were mounted on slides with ProLong™ Gold Antifade Mountant (Catalog # P10144, ThermoFisher Scientific) after washing with PBT. Primary antibodies used in this study include chicken anti-GFP (Catalog #GFP-1020, Aves Labs, Tigard, OR; 1:500–1000), rabbit anti-DesRed (Catalog #632392, Takara Bio USA, Inc., Mountain View, CA; 1:250), rabbit anti-Dpn (1:500) (a gift from Dr. Y.N. Jan) (Bier et al. 1992), guinea pig anti-Ase (a gift from Dr. Y.N. Jan, 1:5000) (Brand et al. 1993), rabbit anti-Miranda (a gift from Dr. Y.N. Jan, 1:1000) (REF), mouse anti-aPKC (Catalog #sc-17781, Santa Cruz Biotechnology, Inc., Dalla, TX, 1:500), rabbit anti-Nito (a gift from N. Perrimon, 1:500) (Yan and Perrimon 2015), mouse anti-V5 mAb (Catalog #R960-25, 1:500 ThermoFisher Scientific) Secondary antibodies conjugated to Daylight 405 (1:300), Daylight 488 (1:100), Cy3 (1:300), Rhodamine Red-X (1:300), Daylight 647 (1:300) or Cy5 (1:300) are from Jackson ImmunoResearch (West Grove, Pennsylvania). Images were collected using a Carl Zeiss LSM780 confocal microscopy and processed with Adobe Photoshop. Student’s T-test was used for statistical analyses.

## Acknowledgments

We thank Drs. N. Perrimon, Y.N. Jan, and C.-Y. Lee for antibodies and fly lines; the Bloomington *Drosophila* Stock Center and the TRiP at Harvard Medical School (NIH/NIGMS R01-GM084947) for providing transgenic RNAi fly stocks used in this study; Addgene for providing plasmids; members of the Pignoni and Lin labs for thoughtful discussion; Chen R. for providing images of *erm-GAL4* expression at early larval stages; Best Gene, Inc for generating transgenic flies; Neuroscience Microscopy Core at Upstate Medical University for providing Zeiss LSM 780 confocal microscopy. This work was supported by the National Institute of Neurological Disorders and Stroke of the National Institutes of Health under Award Number R01NS085232 (S.Z.).

## Author Contributions

Conceptualization, X.L., W.L, M.C., and S.Z.; Methodology, X.L., W.L, M.C., X.D., and S.Z.; Investigation, X.L., W.L, M.C., X.D., and S.Z.; Writing-Original Draft, X.L. W.L. and S.Z, Writing-Review & Editing- X.L., W.L, M.C., and S.Z.; Supervision, S.Z.; Funding, S.Z.

## REFERENCES

Almeida MS, Bray SJ. 2005. Regulation of post-embryonic neuroblasts by Drosophila Grainyhead. Mech Dev 122: 1282–1293.

Andriatsilavo M, Stefanutti M, Siudeja K, Perdigoto CN, Boumard B, Gervais L, Gillet-Markowska A, Al Zouabi L, Schweisguth F, Bardin AJ. 2018. Spen limits intestinal stem cell self-renewal. PLoS Genet 14: e1007773.

Arieti F, Gabus C, Tambalo M, Huet T, Round A, Thore S. 2014. The crystal structure of the Split End protein SHARP adds a new layer of complexity to proteins containing RNA recognition motifs. Nucleic Acids Res 42: 6742–6752.

Artavanis-Tsakonas S, Rand MD, Lake RJ. 1999. Notch signaling: cell fate control and signal integration in development. Science 284: 770–776.

Barolo S, Stone T, Bang AG, Posakony JW. 2002. Default repression and Notch signaling: Hairless acts as an adaptor to recruit the corepressors Groucho and dCtBP to Suppressor of Hairless. Genes Dev 16: 1964–1976.

Bello BC, Izergina N, Caussinus E, Reichert H. 2008. Amplification of neural stem cell proliferation by intermediate progenitor cells in Drosophila brain development. Neural Dev 3: 5.

Berdnik D, Torok T, Gonzalez-Gaitan M, Knoblich JA. 2002. The endocytic protein alpha-Adaptin is required for numb-mediated asymmetric cell division in Drosophila. Dev Cell 3: 221–231.

Bhat KM. 2014. Notch signaling acts before cell division to promote asymmetric cleavage and cell fate of neural precursor cells. Sci Signal 7: ra101.

Bier E, Vaessin H, Younger-Shepherd S, Jan LY, Jan YN. 1992. deadpan, an essential pan-neural gene in Drosophila, encodes a helix-loop-helix protein similar to the hairy gene product. Genes Dev 6: 2137–2151.

Bonev B, Stanley P, Papalopulu N. 2012. MicroRNA-9 Modulates Hes1 ultradian oscillations by forming a double-negative feedback loop. Cell Rep 2: 10–18.

Boone JQ, Doe CQ. 2008. Identification of Drosophila type II neuroblast lineages containing transit amplifying ganglion mother cells. Dev Neurobiol 68: 1185–1195.

Bowman SK, Rolland V, Betschinger J, Kinsey KA, Emery G, Knoblich JA. 2008. The tumor suppressors Brat and Numb regulate transit-amplifying neuroblast lineages in Drosophila. Dev Cell 14: 535–546.

Bowness JS, Nesterova TB, Wei G, Rodermund L, Almeida M, Coker H, Carter EJ, Kadaster A, Brockdorff N. 2022. Xist-mediated silencing requires additive functions of SPEN and Polycomb together with differentiation-dependent recruitment of SmcHD1. Cell Rep 39: 110830.

Brand M, Jarman AP, Jan LY, Jan YN. 1993. asense is a Drosophila neural precursor gene and is capable of initiating sense organ formation. Development 119: 1–17.

Bray SJ. 2006. Notch signalling: a simple pathway becomes complex. Nat Rev Mol Cell Biol 7: 678–689.

Button AC, Hall SD, Ashley EL, McHugh CA. 2024. Dissection of protein and RNA regions required for SPEN binding to XIST A-repeat RNA. RNA 30: 240–255.

Capobianco AJ, Zagouras P, Blaumueller CM, Artavanis-Tsakonas S, Bishop JM. 1997. Neoplastic transformation by truncated alleles of human NOTCH1/TAN1 and NOTCH2. Mol Cell Biol 17: 6265–6273.

Carter AC, Xu J, Nakamoto MY, Wei Y, Zarnegar BJ, Shi Q, Broughton JP, Ransom RC, Salhotra A, Nagaraja SD et al. 2020. Spen links RNA-mediated endogenous retrovirus silencing and X chromosome inactivation. Elife 9: e54508

Ceci M, Fazi F, Romano N. 2021. The role of RNA-binding and ribosomal proteins as specific RNA translation regulators in cellular differentiation and carcinogenesis. Biochim Biophys Acta Mol Basis Dis 1867: 166046.

Chang JL, Lin HV, Blauwkamp TA, Cadigan KM. 2008. Spenito and Split ends act redundantly to promote Wingless signaling. Dev Biol 314: 100–111.

Chen F, Rebay I. 2000. split ends, a new component of the Drosophila EGF receptor pathway, regulates development of midline glial cells. Curr Biol 10: 943–946.

Chen R, Deng X, Zhu S. 2022. The Ets protein Pointed P1 represses Asense expression in type II neuroblasts by activating Tailless. PLoS Genet 18: e1009928.

Chen R, Hou Y, Connell M, Zhu S. 2021. Homeodomain protein Six4 prevents the generation of supernumerary Drosophila type II neuroblasts and premature differentiation of intermediate neural progenitors. PLoS Genet 17: e1009371.

Choksi SP, Southall TD, Bossing T, Edoff K, de Wit E, Fischer BE, van Steensel B, Micklem G, Brand AH. 2006. Prospero acts as a binary switch between self-renewal and differentiation in Drosophila neural stem cells. Dev Cell 11: 775–789.

Chu C, Zhang QC, da Rocha ST, Flynn RA, Bharadwaj M, Calabrese JM, Magnuson T, Heard E, Chang HY. 2015. Systematic discovery of Xist RNA binding proteins. Cell 161: 404–416.

Connell M, Xie Y, Deng X, Chen R, Zhu S. 2024. Kin17 regulates proper cortical localization of Miranda in Drosophila neuroblasts by regulating Flfl expression. Cell Rep 43: 113823.

Couturier L, Vodovar N, Schweisguth F. 2012. Endocytosis by Numb breaks Notch symmetry at cytokinesis. Nat Cell Biol 14: 131–139.

de la Pompa JL, Wakeham A, Correia KM, Samper E, Brown S, Aguilera RJ, Nakano T, Honjo T, Mak TW, Rossant J et al. 1997. Conservation of the Notch signalling pathway in mammalian neurogenesis. Development 124: 1139–1148.

Dirks RM, Pierce NA. 2004. Triggered amplification by hybridization chain reaction. Proc Natl Acad Sci U S A 101: 15275–15278.

Dominissini D, Moshitch-Moshkovitz S, Schwartz S, Salmon-Divon M, Ungar L, Osenberg S, Cesarkas K, Jacob-Hirsch J, Amariglio N, Kupiec M et al. 2012. Topology of the human and mouse m6A RNA methylomes revealed by m6A-seq. Nature 485: 201–206.

Dong Z, Yang N, Yeo SY, Chitnis A, Guo S. 2012. Intralineage directional Notch signaling regulates self-renewal and differentiation of asymmetrically dividing radial glia. Neuron 74: 65–78.

Doroquez DB, Orr-Weaver TL, Rebay I. 2007. Split ends antagonizes the Notch and potentiates the EGFR signaling pathways during Drosophila eye development. Mech Dev 124: 792–806.

Eroglu E, Burkard TR, Jiang Y, Saini N, Homem CCF, Reichert H, Knoblich JA. 2014. SWI/SNF complex prevents lineage reversion and induces temporal patterning in neural stem cells. Cell 156: 1259–1273.

Fre S, Huyghe M, Mourikis P, Robine S, Louvard D, Artavanis-Tsakonas S. 2005. Notch signals control the fate of immature progenitor cells in the intestine. Nature 435: 964–968.

Frierson HF, Jr., Moskaluk CA. 2013. Mutation signature of adenoid cystic carcinoma: evidence for transcriptional and epigenetic reprogramming. J Clin Invest 123: 2783–2785.

Girard V, Goubard V, Querenet M, Seugnet L, Pays L, Nataf S, Dufourd E, Cluet D, Mollereau B, Davoust N. 2020. Spen modulates lipid droplet content in adult Drosophila glial cells and protects against paraquat toxicity. Sci Rep 10: 20023.

Glisovic T, Bachorik JL, Yong J, Dreyfuss G. 2008. RNA-binding proteins and post-transcriptional gene regulation. FEBS Lett 582: 1977–1986.

Gratz SJ, Rubinstein CD, Harrison MM, Wildonger J, O’Connor-Giles KM. 2015. CRISPR-Cas9 Genome Editing in Drosophila. Curr Protoc Mol Biol 111: 31.2.1-31.2.20.

Guo M, Jan LY, Jan YN. 1996. Control of daughter cell fates during asymmetric division: interaction of Numb and Notch. Neuron 17: 27–41.

Hakes AE, Brand AH. 2020. Tailless/TLX reverts intermediate neural progenitors to stem cells driving tumourigenesis via repression of asense/ASCL1. Elife 9: e53377

Hardy EC, Balcerowicz M. 2024. Untranslated yet indispensable-UTRs act as key regulators in the environmental control of gene expression. J Exp Bot 75: 4314–4331.

Henrique D, Hirsinger E, Adam J, Le Roux I, Pourquie O, Ish-Horowicz D, Lewis J. 1997. Maintenance of neuroepithelial progenitor cells by Delta-Notch signalling in the embryonic chick retina. Curr Biol 7: 661–670.

Hiriart E, Gruffat H, Buisson M, Mikaelian I, Keppler S, Meresse P, Mercher T, Bernard OA, Sergeant A, Manet E. 2005. Interaction of the Epstein-Barr virus mRNA export factor EB2 with human Spen proteins SHARP, OTT1, and a novel member of the family, OTT3, links Spen proteins with splicing regulation and mRNA export. J Biol Chem 280: 36935–36945.

Hitoshi S, Alexson T, Tropepe V, Donoviel D, Elia AJ, Nye JS, Conlon RA, Mak TW, Bernstein A, van der Kooy D. 2002. Notch pathway molecules are essential for the maintenance, but not the generation, of mammalian neural stem cells. Genes Dev 16: 846–858.

Janssens DH, Hamm DC, Anhezini L, Xiao Q, Siller KH, Siegrist SE, Harrison MM, Lee CY. 2017. An Hdac1/Rpd3-Poised Circuit Balances Continual Self-Renewal and Rapid Restriction of Developmental Potential during Asymmetric Stem Cell Division. Dev Cell 40: 367–380 e367.

Janssens DH, Komori H, Grbac D, Chen K, Koe CT, Wang H, Lee CY. 2014. Earmuff restricts progenitor cell potential by attenuating the competence to respond to self-renewal factors. Development 141: 1036–1046.

Jemc J, Rebay I. 2006. Characterization of the split ends-like gene spenito reveals functional antagonism between SPOC family members during Drosophila eye development. Genetics 173: 279–286.

Jin LH, Choi JK, Kim B, Cho HS, Kim J, Kim-Ha J, Kim YJ. 2009. Requirement of Split ends for epigenetic regulation of Notch signal-dependent genes during infection-induced hemocyte differentiation. Mol Cell Biol 29: 1515–1525.

Jinek M, Chylinski K, Fonfara I, Hauer M, Doudna JA, Charpentier E. 2012. A programmable dual-RNA-guided DNA endonuclease in adaptive bacterial immunity. Science 337: 816–821.

Kan L, Grozhik AV, Vedanayagam J, Patil DP, Pang N, Lim KS, Huang YC, Joseph B, Lin CJ, Despic V et al. 2017. The m(6)A pathway facilitates sex determination in Drosophila. Nat Commun 8: 15737.

Knuckles P, Lence T, Haussmann IU, Jacob D, Kreim N, Carl SH, Masiello I, Hares T, Villasenor R, Hess D et al. 2018. Zc3h13/Flacc is required for adenosine methylation by bridging the mRNA-binding factor Rbm15/Spenito to the m(6)A machinery component Wtap/Fl(2)d. Genes Dev 32: 415–429.

Koe CT, Li S, Rossi F, Wong JJ, Wang Y, Zhang Z, Chen K, Aw SS, Richardson HE, Robson P et al. 2014. The Brm-HDAC3-Erm repressor complex suppresses dedifferentiation in Drosophila type II neuroblast lineages. Elife 3: e01906.

Kuang B, Wu SC, Shin Y, Luo L, Kolodziej P. 2000. split ends encodes large nuclear proteins that regulate neuronal cell fate and axon extension in the Drosophila embryo. Development 127: 1517–1529.

Kuroda K, Han H, Tani S, Tanigaki K, Tun T, Furukawa T, Taniguchi Y, Kurooka H, Hamada Y, Toyokuni S et al. 2003. Regulation of marginal zone B cell development by MINT, a suppressor of Notch/RBP-J signaling pathway. Immunity 18: 301–312.

Lai EC, Burks C, Posakony JW. 1998. The K box, a conserved 3’ UTR sequence motif, negatively regulates accumulation of enhancer of split complex transcripts. Development 125: 4077–4088.

Lai EC, Posakony JW. 1997. The Bearded box, a novel 3’ UTR sequence motif, mediates negative post-transcriptional regulation of Bearded and Enhancer of split Complex gene expression. Development 124: 4847–4856.

Lai EC, Tam B, Rubin GM. 2005. Pervasive regulation of Drosophila Notch target genes by GY-box-, Brd-box-, and K-box-class microRNAs. Genes Dev 19: 1067–1080.

Le Borgne R, Schweisguth F. 2003. Unequal segregation of Neuralized biases Notch activation during asymmetric cell division. Dev Cell 5: 139–148.

Lence T, Akhtar J, Bayer M, Schmid K, Spindler L, Ho CH, Kreim N, Andrade-Navarro MA, Poeck B, Helm M et al. 2016. m(6)A modulates neuronal functions and sex determination in Drosophila. Nature 540: 242–247.

Li B, Wong C, Gao SM, Zhang R, Sun R, Li Y, Song Y. 2018. The retromer complex safeguards against neural progenitor-derived tumorigenesis by regulating Notch receptor trafficking. Elife 7: e38181

Li L, Vaessin H. 2000. Pan-neural Prospero terminates cell proliferation during Drosophila neurogenesis. Genes Dev 14: 147–151.

Li X, Chen R, Zhu S. 2017. bHLH-O proteins balance the self-renewal and differentiation of Drosophila neural stem cells by regulating Earmuff expression. Dev Biol 431: 239–251.

Li X, Xie Y, Zhu S. 2016. Notch maintains Drosophila type II neuroblasts by suppressing expression of the Fez transcription factor Earmuff. Development 143: 2511–2521.

Lindtner S, Zolotukhin AS, Uranishi H, Bear J, Kulkarni V, Smulevitch S, Samiotaki M, Panayotou G, Felber BK, Pavlakis GN. 2006. RNA-binding motif protein 15 binds to the RNA transport element RTE and provides a direct link to the NXF1 export pathway. J Biol Chem 281: 36915–36928.

Liu K, Shen D, Shen J, Gao SM, Li B, Wong C, Feng W, Song Y. 2017. The Super Elongation Complex Drives Neural Stem Cell Fate Commitment. Dev Cell 40: 537–551 e536.

Liu Q, Wang XY, Qin YY, Yan XL, Chen HM, Huang QD, Chen JK, Zheng JM. 2018. SPOCD1 promotes the proliferation and metastasis of glioma cells by up-regulating PTX3. Am J Cancer Res 8: 624–635.

Luo L, Liao YJ, Jan LY, Jan YN. 1994. Distinct morphogenetic functions of similar small GTPases: Drosophila Drac1 is involved in axonal outgrowth and myoblast fusion. Genes Dev 8: 1787–1802.

Ma X, Renda MJ, Wang L, Cheng EC, Niu C, Morris SW, Chi AS, Krause DS. 2007. Rbm15 modulates Notch-induced transcriptional activation and affects myeloid differentiation. Mol Cell Biol 27: 3056–3064.

Ma Z, Morris SW, Valentine V, Li M, Herbrick JA, Cui X, Bouman D, Li Y, Mehta PK, Nizetic D et al. 2001. Fusion of two novel genes, RBM15 and MKL1, in the t(1;22)(p13;q13) of acute megakaryoblastic leukemia. Nat Genet 28: 220–221.

Maier D. 2019. The evolution of transcriptional repressors in the Notch signaling pathway: a computational analysis. Hereditas 156: 5.

McHugh CA, Chen CK, Chow A, Surka CF, Tran C, McDonel P, Pandya-Jones A, Blanco M, Burghard C, Moradian A et al. 2015. The Xist lncRNA interacts directly with SHARP to silence transcription through HDAC3. Nature 521: 232–236.

McLaren M, Butts J. 2025. Notch signaling in neurogenesis. Development 152.

Meyer KD, Saletore Y, Zumbo P, Elemento O, Mason CE, Jaffrey SR. 2012. Comprehensive analysis of mRNA methylation reveals enrichment in 3’ UTRs and near stop codons. Cell 149: 1635–1646.

Mignone F, Gissi C, Liuni S, Pesole G. 2002. Untranslated regions of mRNAs. Genome Biol 3: REVIEWS0004.

Morel V, Lecourtois M, Massiani O, Maier D, Preiss A, Schweisguth F. 2001. Transcriptional repression by suppressor of hairless involves the binding of a hairless-dCtBP complex in Drosophila. Curr Biol 11: 789–792.

Murakami S, Jaffrey SR. 2022. Hidden codes in mRNA: Control of gene expression by m(6)A. Mol Cell 82: 2236–2251.

Naderi A. 2018. SRARP and HSPB7 are epigenetically regulated gene pairs that function as tumor suppressors and predict clinical outcome in malignancies. Mol Oncol 12: 724–755.

Oswald F, Kostezka U, Astrahantseff K, Bourteele S, Dillinger K, Zechner U, Ludwig L, Wilda M, Hameister H, Knochel W et al. 2002. SHARP is a novel component of the Notch/RBP-Jkappa signalling pathway. EMBO J 21: 5417–5426.

Oswald F, Winkler M, Cao Y, Astrahantseff K, Bourteele S, Knochel W, Borggrefe T. 2005. RBP-Jkappa/SHARP recruits CtIP/CtBP corepressors to silence Notch target genes. Mol Cell Biol 25: 10379–10390.

Patil DP, Chen CK, Pickering BF, Chow A, Jackson C, Guttman M, Jaffrey SR. 2016. m(6)A RNA methylation promotes XIST-mediated transcriptional repression. Nature 537: 369–373.

Pfeiffer BD, Jenett A, Hammonds AS, Ngo TT, Misra S, Murphy C, Scully A, Carlson JW, Wan KH, Laverty TR et al. 2008. Tools for neuroanatomy and neurogenetics in Drosophila. Proc Natl Acad Sci U S A 105: 9715–9720.

Querenet M, Goubard V, Chatelain G, Davoust N, Mollereau B. 2015. Spen is required for pigment cell survival during pupal development in Drosophila. Dev Biol 402: 208–215.

Razumova E, Makariuk A, Dontsova O, Shepelev N, Rubtsova M. 2025. Structural Features of 5’ Untranslated Region in Translational Control of Eukaryotes. Int J Mol Sci 26.

Rives-Quinto N, Komori H, Ostgaard CM, Janssens DH, Kondo S, Dai Q, Moore AW, Lee CY. 2020. Sequential activation of transcriptional repressors promotes progenitor commitment by silencing stem cell identity genes. Elife 9: e56187

Robert-Finestra T, Tan BF, Mira-Bontenbal H, Timmers E, Gontan C, Merzouk S, Giaimo BD, Dossin F, van IWFJ, Martens JWM et al. 2021. SPEN is required for Xist upregulation during initiation of X chromosome inactivation. Nat Commun 12: 7000.

Rodermund L, Coker H, Oldenkamp R, Wei G, Bowness J, Rajkumar B, Nesterova T, Susano Pinto DM, Schermelleh L, Brockdorff N. 2021. Time-resolved structured illumination microscopy reveals key principles of Xist RNA spreading. Science 372.

Rubin GM, Hong L, Brokstein P, Evans-Holm M, Frise E, Stapleton M, Harvey DA. 2000. A Drosophila complementary DNA resource. Science 287: 2222–2224.

Ryczek N, Lys A, Makalowska I. 2023. The Functional Meaning of 5’UTR in Protein-Coding Genes. Int J Mol Sci 24: 2976

San-Juan BP, Baonza A. 2011. The bHLH factor deadpan is a direct target of Notch signaling and regulates neuroblast self-renewal in Drosophila. Dev Biol 352: 70–82.

Sanchez-Pulido L, Rojas AM, van Wely KH, Martinez AC, Valencia A. 2004. SPOC: a widely distributed domain associated with cancer, apoptosis and transcription. BMC Bioinformatics 5: 91.

Song Y, Lu B. 2012. Interaction of Notch signaling modulator Numb with alpha-Adaptin regulates endocytosis of Notch pathway components and cell fate determination of neural stem cells. J Biol Chem 287: 17716–17728.

Spana EP, Doe CQ. 1996. Numb antagonizes Notch signaling to specify sibling neuron cell fates. Neuron 17: 21–26.

Stockhausen MT, Kristoffersen K, Poulsen HS. 2010. The functional role of Notch signaling in human gliomas. Neuro Oncol 12: 199–211.

Su S, Li S, Deng T, Gao M, Yin Y, Wu B, Peng C, Liu J, Ma J, Zhang K. 2022. Cryo-EM structures of human m(6)A writer complexes. Cell Res 32: 982–994.

Tsuneoka Y, Funato H. 2020. Modified in situ Hybridization Chain Reaction Using Short Hairpin DNAs. Front Mol Neurosci 13: 75.

Uranishi H, Zolotukhin AS, Lindtner S, Warming S, Zhang GM, Bear J, Copeland NG, Jenkins NA, Pavlakis GN, Felber BK. 2009. The RNA-binding motif protein 15B (RBM15B/OTT3) acts as cofactor of the nuclear export receptor NXF1. J Biol Chem 284: 26106–26116.

van der Heijden AG, Mengual L, Lozano JJ, Ingelmo-Torres M, Ribal MJ, Fernandez PL, Oosterwijk E, Schalken JA, Alcaraz A, Witjes JA. 2016. A five-gene expression signature to predict progression in T1G3 bladder cancer. Eur J Cancer 64: 127–136.

Wang H, Somers GW, Bashirullah A, Heberlein U, Yu F, Chia W. 2006. Aurora-A acts as a tumor suppressor and regulates self-renewal of Drosophila neuroblasts. Genes Dev 20: 3453–3463.

Weng M, Golden KL, Lee CY. 2010. dFezf/Earmuff maintains the restricted developmental potential of intermediate neural progenitors in Drosophila. Dev Cell 18: 126–135.

Wiellette EL, Harding KW, Mace KA, Ronshaugen MR, Wang FY, McGinnis W. 1999. spen encodes an RNP motif protein that interacts with Hox pathways to repress the development of head-like sclerites in the Drosophila trunk. Development 126: 5373–5385.

Yan D, Perrimon N. 2015. spenito is required for sex determination in Drosophila melanogaster. Proc Natl Acad Sci U S A 112: 11606–11611.

Zaccara S, Ries RJ, Jaffrey SR. 2019. Reading, writing and erasing mRNA methylation. Nat Rev Mol Cell Biol 20: 608–624.

Zacharioudaki E, Magadi SS, Delidakis C. 2012. bHLH-O proteins are crucial for Drosophila neuroblast self-renewal and mediate Notch-induced overproliferation. Development 139: 1258–1269.

Zhang L, Tran NT, Su H, Wang R, Lu Y, Tang H, Aoyagi S, Guo A, Khodadadi-Jamayran A, Zhou D et al. 2015. Cross-talk between PRMT1-mediated methylation and ubiquitylation on RBM15 controls RNA splicing. Elife 4: 07938

Zhu S, Barshow S, Wildonger J, Jan LY, Jan YN. 2011. Ets transcription factor Pointed promotes the generation of intermediate neural progenitors in Drosophila larval brains. Proc Natl Acad Sci U S A 108: 20615–20620.

Zhu S, Wildonger J, Barshow S, Younger S, Huang Y, Lee T. 2012. The bHLH repressor Deadpan regulates the self-renewal and specification of Drosophila larval neural stem cells independently of Notch. PLoS One 7: e46724.

